# The asynchrony in the exit from naive pluripotency cannot be explained by differences in the cell cycle phase

**DOI:** 10.1101/2023.09.15.557731

**Authors:** Swathi Jayaram, Merrit Romeike, Christa Buecker

## Abstract

Development is characterized by consecutive cell state transitions that build on each other and ultimately lead to the generation of the numerous different cell types found in the organism. During each of these transitions, cells change their gene expression profiles and take on new identities. Cell state transitions have to be tightly coordinated with proliferation to ensure simultaneous growth and differentiation. The exit from naive pluripotency is an ideal model system for studying the temporal coordination of proliferation and differentiation. Individual cells initiate differentiation earlier compared to others, thereby leading to an asynchronous exit from naive pluripotency. One of the major differences among the cells of the starting population of mouse embryonic stem cells (mESCs) is the cell cycle status, and could therefore be an underlying cause of the differences in the onset of the exit from naive pluripotency. However, through comprehensive analysis including single cell RNA sequencing (scRNA-seq), cell cycle synchronization, and perturbation experiments, we demonstrate here that the cell cycle phase at the initiation of differentiation does not influence the timing of the exit from naive pluripotency.

## INTRODUCTION

Consecutive cell state transitions during development eventually lead to the different cellular states that together make up the whole organism. With each cell state transition, a new transcriptional program is established, setting the stage for the next cell state transition. One ideal model system to study such cell state transitions in a highly controlled manner is the exit from naive pluripotency^1–3^: murine embryonic stem cells (mESCs) represent the so-called naive pluripotent stem cell state and are cultured under defined conditions through stimulation with Leukemia Inhibitory Factor (LIF) and inhibition of differentiation signals through the usage of two inhibitors (2i) against MEK/ERK signalling and GSK3b, which in turn stimulates Wnt activity^4–6^. When 2i and LIF (2iLIF) are removed and replaced instead by stimulation with bFGF, the cells exit the naive state of pluripotency and initiate differentiation into the next developmental state, often referred to as formative pluripotency^7–10^. Within 24-36hrs the cells downregulate the naive pluripotent gene expression profile and activate a novel gene expression profile^11–15^, thereby committing to differentiation irreversibly.

Individual mESCs do not respond to the differentiation cue simultaneously: some cells initiate differentiation early, while others lag behind and only change their transcriptional profile hours later^16,17^. It is an outstanding question, what determines when a cell initiates differentiation and what is the underlying cause of the asynchrony in differentiation. 2iLIF cultured mESCs are more homogeneous compared to other culture conditions such as serum/LIF^18–21^, and the remaining variability is predominantly caused by differences in the expression of genes related to the cell cycle^18^. Conversely, mESCs cultured in serum/LIF conditions show heterogeneity in genes related to developmentally distinct cellular states^18^.

During development, cells do not just change their transcriptional profiles, they also proliferate. Both processes have to be coordinated to ensure robust development. Beyond simple coordination of the two processes, it was suggested that the cell cycle phase of a given cell determines its propensity to differentiate^22–25^. For example, in human embryonic stem cells (hESCs), the G1 phase offers a window of opportunity for cells to initiate differentiation through early G1 specific SMAD signalling activity facilitated by cyclin D1-D3^26^. Furthermore, deposition of bivalent histone marks on developmental genes during G1 phase through CDK2 activity leads to changes in enhancer-promoter contacts and subsequent upregulation of developmental genes^27,28^.

In mESCs, mitotic exit is accompanied by fast reactivation of self-renewal genes bookmarked by H3K27ac, rapid enhancer reactivation, and re-establishment of chromatin architecture^29^.

Differential expression of PRC2 variant proteins EPOP and EloB across the cell cycle has been linked to the upregulation of lineage-specific markers in G1 phase^30^. G1 phase synchronization experiments in mESCs, followed by differentiation and cell state analysis using RT-qPCR suggested that inhibition of mitosis and S phase leads to impaired exit from naive pluripotency^31^. This study however was limited to cells sorted from the G1 cell cycle state and did not test how other cell cycle phases at the time point of differentiation initiation influence the decision to exit from naive pluripotency. Live cell imaging of the Rex1-GFPd2 reporter system^32^ was used to address this question, but two studies reached opposing conclusions. In this system, the expression of a destabilized GFP variant is under the control of the naive specific promoter of *Rex1*, therefore downregulation of GFP indicates the exit from naive pluripotency. Chaigne A. *et al.* reported that mESCs exit from naive pluripotency only after one cell cycle^33^, while Strawbridge S. *et al*. reported that mESCs pass through variable lag phases before exiting from naive pluripotency and that the propensity to differentiate is not linked to a specific cell cycle phase^17^. Taken together, the potential connection between the cell cycle phase and differentiation initiation is still not fully understood.

In this study, we set out to investigate if the cell cycle phase at the initiation of differentiation is the underlying source of asynchrony in the exit from naive pluripotency. As mESCs are released from the self-renewal conditions, irrespective of the differentiation status, they have equal representation of all the cell cycle phases. This indicates that cell cycle and cell state are not tightly linked. Furthermore, cell cycle synchronization followed by differentiation revealed that most cell state specific markers do not show a cell cycle dependency in expression. Moreover, cell cycle synchronization did not lead to a synchronized exit from naive pluripotency, indicating that the cell cycle at the initiation of differentiation does not influence the timing of this transition. Finally, mutations causing a differentiation delay do not exhibit any change in the cell cycle profile compared to the WT. All of these individual results together strongly suggest that the cell cycle is not a major cause of the asynchrony in the exit from naive pluripotency.

## RESULTS

### scRNA-seq expression analysis to interrogate intermediate time points of the exit from naive pluripotency

The exit from naive pluripotency is an ideal model system to study the link between proliferation and cell state transitions: the starting population is highly homogenous and the main differences among the cells have been attributed to the cell cycle^18^. To analyse the gene expression changes during the exit from naive pluripotency, we performed single cell RNA-seq (scRNA-seq) at selected time points upon initiation of differentiation. While naive and formative pluripotency have distinct transcriptional profiles^11^, they still represent closely connected cell states. To perform scRNA-seq with an unconfounded experimental setup and no necessity for batch correction, we staggered the onset of differentiation and harvested and processed cells in parallel (Fig S1A). In addition, we pooled cells from all time points using the MULTI-seq pooling approach^34^, further reducing the introduction of technical confounders like encapsulation reaction or sequencing depth. We validated labelling efficiency and label retention in our system by incorporating fluorescently labelled DNA barcodes into the membrane of cells followed by FACS analysis, analogous to the MULTI-seq development. Cells were efficiently labelled with barcodes and clearly separated by FACS profiles (Fig S1B, left panel). Next, we pooled two distinctly labelled cell populations and incubated them together to test if barcodes can be exchanged between cells. Cells clearly clustered into two distinct populations and no double positive cells were detected (Fig S1B, right panel). We concluded that MULTI-seq is an efficient method to multiplex cells throughout our time course.

We pooled equal numbers of cells from 5 time points in differentiation. To capture the imminent response to the onset of differentiation we selected timepoints at 6 h and 12 h after differentiation initiation, followed by cells taken at 24 h and 48 h after differentiation initiation. We assigned time point information to the cells. Of note, MULTI-seq introduces a stringent way to identify cell doublets, as droplets with conflicting cell barcode information are identified (Fig S1C). The doublets identified also showed an increased number of detected genes (Fig S1D) and were removed before further analysis. For some cells, no associated timepoint could be identified suggesting that these cells represented a random, unlabelled mixture from all experimental time points (“NaN” in Fig S1C and Fig S1D). We retained these cells in downstream analysis and used them as an internal reference. We first evaluated the expression of naive (*Tbx3, Rex1*) and formative (*Otx2*) marker genes across experimental time points, which showed expected down-and upregulation behaviour, respectively, albeit with marker specific expression levels and kinetics (Fig S1E). Therefore, experimental time points were correctly identified after demultiplexing.

Next, we analysed the scRNA-seq dataset by performing an unsupervised principal component analysis (PCA) based on the top 500 most variable genes within the data. We observed that the clustering does not recapitulate the order of time points analysed with the 0 h misplaced along principal component 1 (PC1) (Fig 1A). Since a link between cell cycle and cell fate has been reported^26,31,33^ we performed *in silico* cell cycle assignment to the scRNA-seq data using Cyclone^35^ to test if cell cycle genes were responsible for the differences in the analysis. Cells in the same cell cycle phase were indeed clustered along PC2 with the G1 phase clustering at the top and the G2 phase at the bottom of the manifold (Fig S2A). We performed linear regression with cell cycle as a blocking factor to test whether this would recapitulate expected differentiation behaviour. Regressing out the cell cycle phase led to the mixing of the cell cycle phases (Fig S2B), however, the distribution of time points did not recapitulate the expected separation of cells based on the known experimental time point. Cells collected at 0 h formed an intermediate state, as observed in PCA and pseudotime alignment of the experimental time points (Fig S2 C, D (pseudotime)). These observations suggest that a transcriptional signature of cycling cells is a different signature than differentiation.

**Fig 1:**
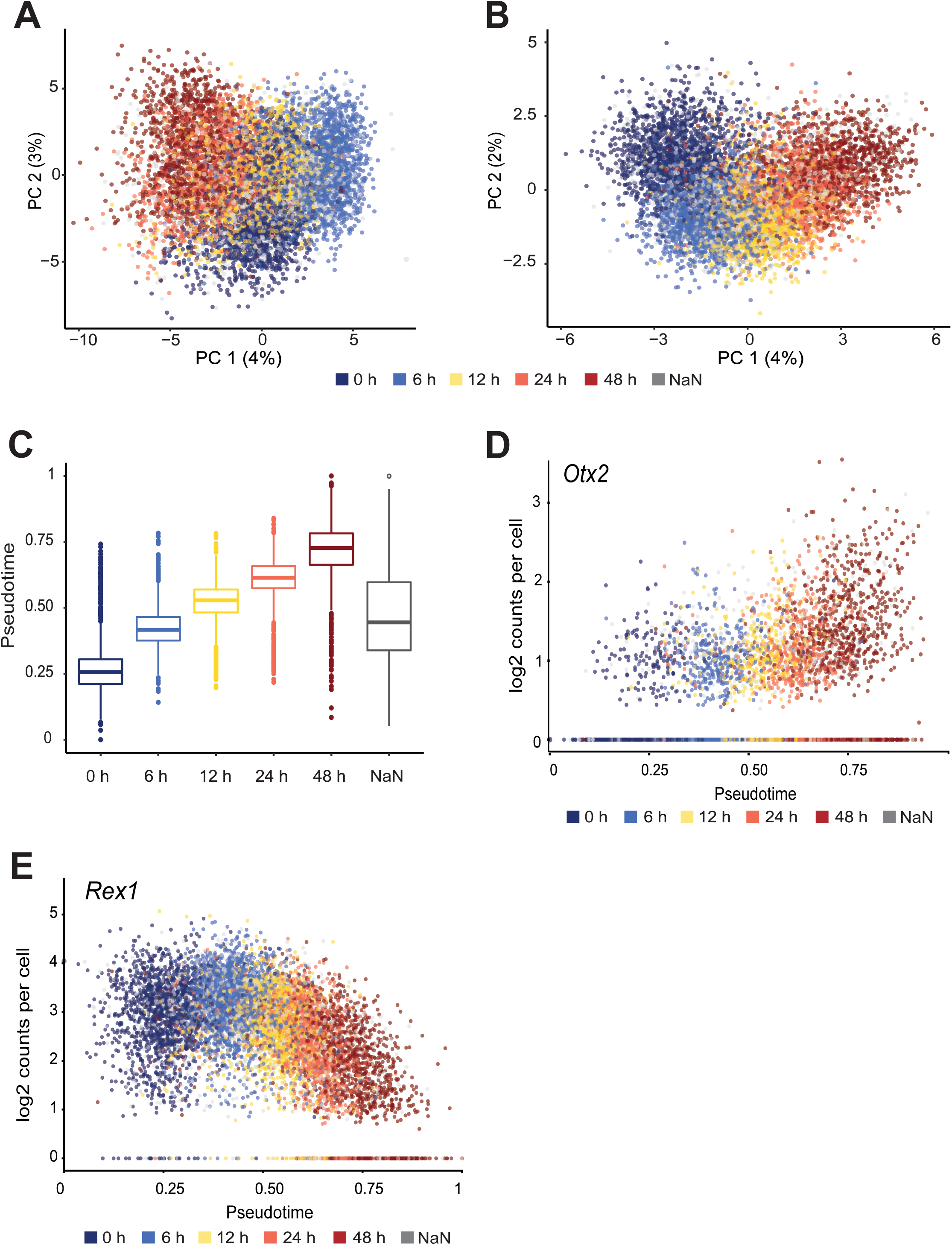
The exit from naive pluripotency is heterogeneous. A. Unsupervised dimension reduction using Principal Component Analysis (PCA) of scRNA-seq data of differentiating mESCs collected at indicated time points (colors) during the exit from naive pluripotency. PCA is based on the top 500 of the most variable genes within the scRNA-seq data B. Supervised dimension reduction using PCA of scRNA-seq data of differentiating mESCs collected at indicated time points (colors) during the exit from naive pluripotency. PCA is based on the top 500 of the differentially expressed genes (DEG) between 0 h and 48 h in bulk RNA seq represented in Fig S2E C. Distribution of pseudotime values of cells collected at an indicated experimental time point, analyzed with supervised dimension reduction using PCA based on top 500 of the differentially expressed genes (DEG) between 0 h and 48 h in bulk RNA seq represent­ed in Fig S2E D. Single cell expression (log2) counts of *Otx2* plotted as a function of pseudotime values with information about the experimental time point indicated in color E. Single cell expression (log2) counts of *Rex1* plotted as a function of pseudotime values with information about the experimental time point indicated in color

### The exit from naive pluripotency is heterogenous

Unsupervised PCA analysis did not recapitulate the progression of known experimental time points. To overcome this limitation, we turned to supervised dimension reduction: first, we performed bulk RNA-seq and selected the top 500 most differentially expressed genes between 0 h and 48 h differentiated cells (Fig S2E) to perform supervised PCA of the scRNA-seq data. Surprisingly there was little overlap between the top 500 variable genes based on scRNA-seq and differentially expressed genes in bulk RNA-seq (Fig S2F), indicating that cell cycle and differentiation are orthogonal processes: major differences in the scRNA-seq data arose due to the cell cycle genes but contributed only minimally to the differentially expressed genes between the differentiated state compared to undifferentiated state. Supervised dimension reduction captured the expected order of time points along the PC1 with 0 h followed by 6 h, 12 h, and up to 48 h (Fig 1B). We assigned pseudotime values to every cell agnostic to the experimental collection time point. Within one collection of time points cells showed a wide spread of pseudotime values (Fig 1C). In other words, some cells harvested from the same time point, and therefore being exposed to the differentiation cue for the same duration, have lower pseudotime values while others have higher pseudotime values. This observation was also recapitulated by the heterogeneity in the upregulation of *Otx2* (Fig 1D) and downregulation of *Rex1* (Fig 1E) and within every time point during the cell fate transition. Taken together, the exit from naive pluripotency is a continuous state transition without discrete intermediate cell states, which exhibits heterogeneity within collection time points.

### Differentiation status differences within a time point are not correlated to the cell cycle

The major differences among mESCs cultured under 2iLIF conditions have been attributed to differences in the cell cycle phase^18^. It is therefore tempting to speculate that the asynchronous exit from naive pluripotency, which leads to the observed heterogeneity, is due to these underlying differences and cells in different phases of the cell cycle react differently to the differentiation cues. During the exit from naive pluripotency, cells do not undergo a cell fate choice but rather follow a single cell state change (Fig 1B) and the evidence for a link between the cell cycle and the asynchronous cell state transition of mESCs remains inconclusive^17,31,33^.

We investigated a possible link between cell cycle and differentiation status by first assigning cell cycle status to each cell in our single cell sequencing data using Cyclone^35^. We reasoned that if the cell cycle phase influences the onset of differentiation and thereby the asynchrony in the exit from naive pluripotency, cells with a higher pseudotime value but collected at the same time point should show enrichment of one of the cell cycle phases. We plotted the cells at different experimental time points along their pseudo time (Fig S2G). Assuming a correlation between cell cycle and differences in differentiation status, splitting cells by cell cycle phase should result in a distinct distribution of pseudotime values. Within one experimental time point, the cell cycle state does not influence pseudotime values (Fig 2B). In addition, we inspected the expression of *Rex1* across all cells within our time course. *Rex1* was downregulated in all the cells irrespective of the cell cycle phase across the time course (Fig 2C). Taken together, the differentiation status of cells from any experimental time point does not show a cell cycle bias.

**Fig 2:**
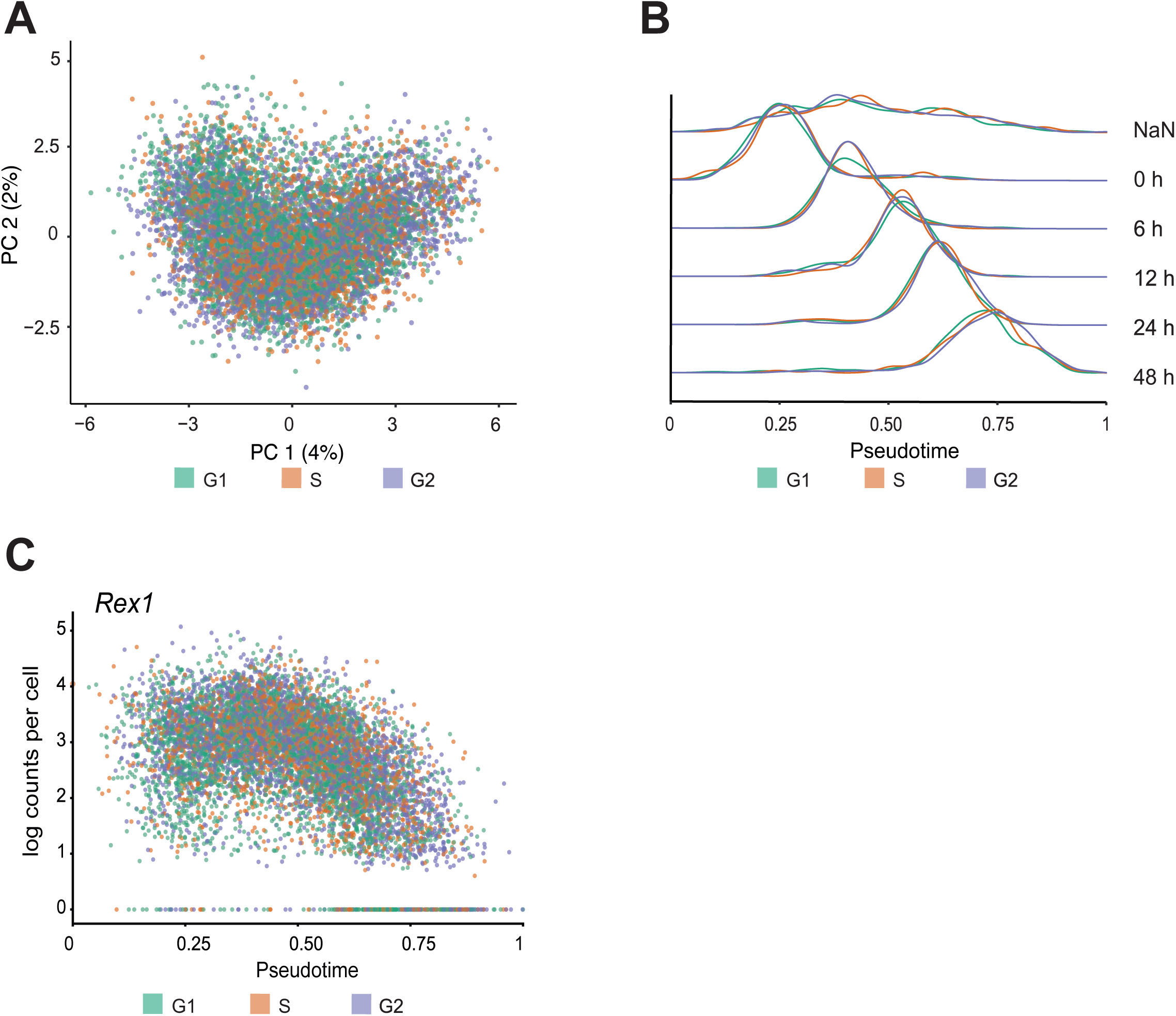
Differentiation status differences within a time point are not correlated to the cell cycle. A. PCA of scRNA-seq data after supervised dimension reduction overlaid with cell cycle phases detected using Cyclone as indicated by the colors (as in Fig 1B) B. Cell cycle assignment using Cyclone to the cells collected at each experimental time point plotted against their pseudotime values (as in Fig S2G) C. Single cell expression (Iog2) counts of *Rex1* plotted as a function of pseudotime values with information about the cell cycle assigned using Cyclone indicated in color (as in Fig 1E)

### The cell cycle phase at the initiation of differentiation does not influence the propensity to differentiate

Cell cycle assignment in the scRNA-seq only represents a snapshot of the time point at analysis but lacks clear information on the cell cycle phase at the initiation of the cell state transition. To overcome this limitation, we performed a cell cycle synchronization experiment to initiate differentiation from a known cell cycle phase and analysed how the cell cycle phase impacts the timing of exit from naive pluripotency. We isolated mESCs based on their DNA content, and thereby cell cycle phases, using Hoechst 33342, a live cell DNA staining dye, through fluorescence activated cell sorting (FACS). After sorting, we immediately initiated differentiation for either 12 h or 16 h. Next, we analysed the differentiation status using real-time quantitative PCR (RT-qPCR) (Fig 3A). We decided to not use the FUCCI system for these experiments since it lacks resolution in separating the S and G2M phases.

**Fig 3:**
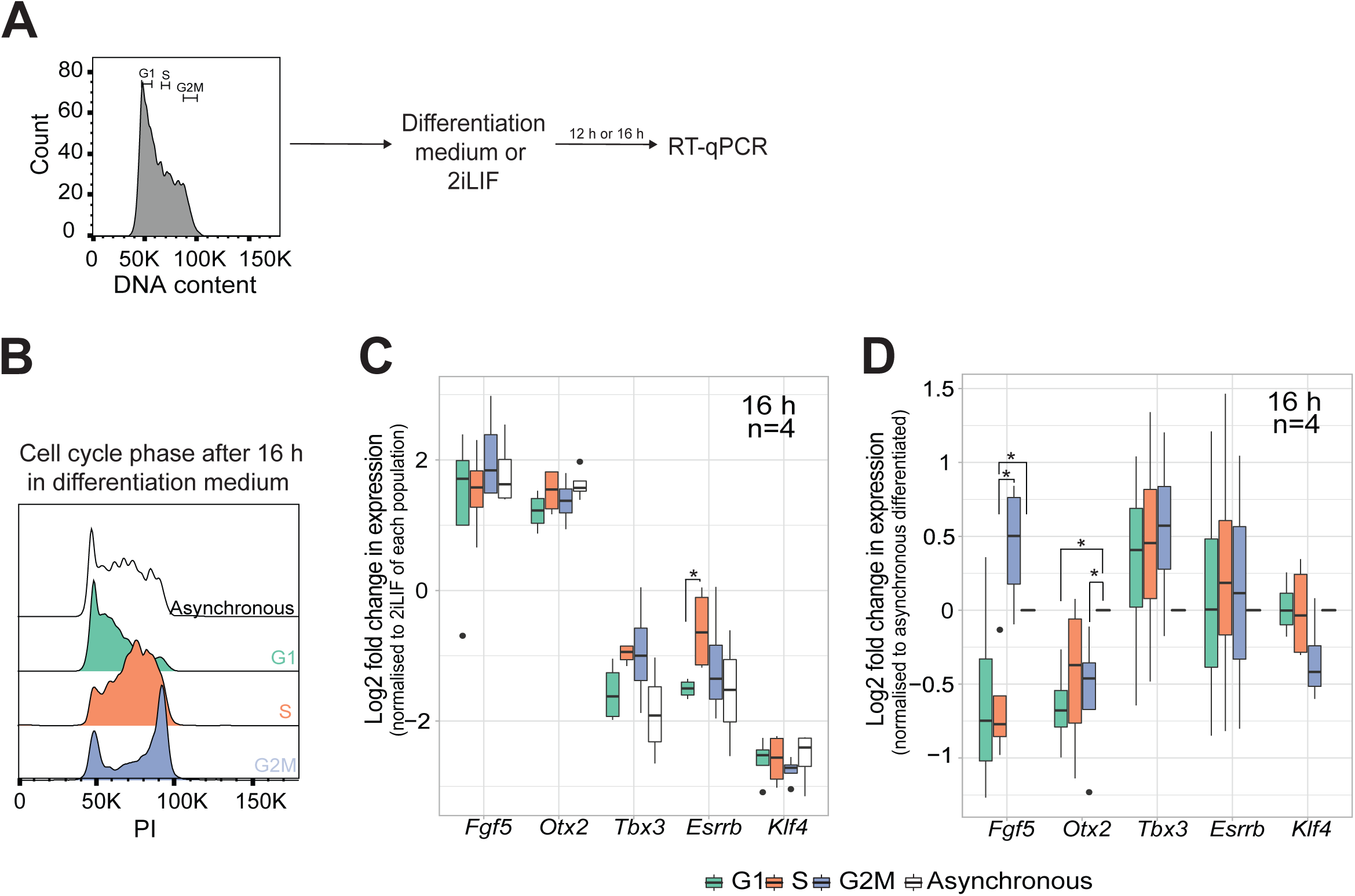
The cell cycle phase at the initiation does not influence the propensity to differentiate. A. Experimental setup of cell cycle isolation and differentiation: Cell cycle status was determined based on DNA content using Hoechst 33342. Cells from either G1, S, or G2M were FACS sorted together with the asyn­chronous population as control and directly plated into the differentiation medium or 2İLIF. After the indicated time point, RNAwas isolated to determine the differentiation status using RT-qPCR B. Analysis of the cell cycle distribution using PI staining, at 16 h post sorting and differentiation of G1, S, G2M, and asynchronous sorted cells C. RT-qPCR of formative *(Fgf5 and Otx2*) and naive markers *(Tbx3, Esrrb, Klf4)* at 16 h post sorting in differentiation medium. Fold changes of gene expression were calculated by normalization to undifferentiated counterparts of each sorted population and Iog2 transformed. Unpaired Wilcoxon test was performed on the Iog2 transformed values, p<0.05 is considered significant, n=4 biological replicates D. RT-qPCR of formative *(Fgf5 and Otx2*) and naive markers *(Tbx3, Esrrb, Klf4)* at 16 h post sorting in differ­entiation medium. Fold changes of gene expression were calculated by normalization to the asynchronous differ­entiated sample and Iog2 transformed. Unpaired Wilcoxon test was performed on the Iog2 transformed values, p<O.05 is considered significant, n=4 biological replicates

Differentiated cells sorted from each cell cycle phase were predominantly found in the same cell cycle phase at 12 h and entered the next cell cycle phase at 16 h, in line with the cell cycle duration of 12 h for mESCs^36^. For example, cells sorted from the G1 phase were found predominantly again in the G1 phase at 12 h and in the G1 and S phase at 16 h. (12 h: Fig S3B right panel; 16 h: Fig 3B). These observations were not limited to differentiation but were also detected in the undifferentiated control cells (12 h: Fig S3B left panel; 16 h: Fig S3A). We analysed the expression levels of different formative (*Fgf5, Otx2*) and naive pluripotency markers (*Esrrb, Tbx3, Klf4*) using RT-qPCR. We first compared the differentiation status of the cells based on the starting cell cycle phase by normalizing the expression to the respective undifferentiated counterparts. At 12 h and 16 h, naive markers were downregulated and formative markers were upregulated to similar levels compared to the asynchronous control. Only *Esrrb* expression was significantly downregulated in cells that were induced to exit from pluripotency from G1 compared to the S phase at 16 h (Fig 3C).

Next, we compared the differentiation status of the cell cycle sorted cells directly to the asynchronous differentiated control. If the cell cycle were a major contributor to the timing of exit from naive pluripotency, then the analysed markers would show significantly changed expression in one of the synchronized populations. At 16 h, none of the naive pluripotency markers showed significant differences to the asynchronous differentiated control, while the analysed formative markers showed cell cycle dependency. However, if it were indeed a cell cycle-based effect then both formative markers should have been upregulated. Instead, we observed that *Otx2* is downregulated in cells irrespective of the starting cell cycle while *Fgf5* is either unchanged or upregulated compared to the asynchronous control (Fig 3D). At 12 h, we observed the same opposing expression of *Fgf5* and *Otx2* compared to asynchronous control. Furthermore, at 12 h the naive marker *Klf4* was upregulated in all the synchronized populations compared to the asynchronous control (Fig S3D).

Taken together, the overall cell state change is not cell cycle dependent compared to the asynchronous population at 12 h or 16 h. *Fgf5* is regulated by the cell cycle, in agreement with a previous report^31^. Therefore, these results indicate that the cell cycle phase at the beginning of differentiation does not influence the timing of the exit from naive pluripotency.

### Cell cycle synchronization does not lead to a more synchronous exit from naive pluripotency

Next, we reasoned that if the cell cycle at the initiation of differentiation were a major source of the asynchrony in the exit from naive pluripotency, then initiating cell state change from a cell cycle synchronized population should lead to a more synchronous transition. In such a case, the naive/formative markers should be expressed more homogeneously throughout the population of differentiating cells. To test this hypothesis, we synchronized cells in mitosis using mitotic shake-off and released them into differentiation for 16 h (Fig 4A). Briefly, mESCs were treated with Nocodazole, a drug that impairs metaphase spindle formation and thereby arrests cells in mitosis. The mitotic cells round up and can be isolated by careful pipetting from the plate (a process called mitotic shake-off) ^29^. The cell cycle purity after shake-off was assessed by Hoechst 33342 staining. The extracted cells are enriched for mitotic cells (Fig 4B) while the DMSO control (i.e., trypsinized DMSO treated cells) (asynchronous) populated all the cell cycle phases. To confirm that the cells released from mitosis re-enter the cell cycle, we labelled newly synthesized DNA with a 30 mins EdU pulse at 3 h post release in an independent experiment (Fig 4C, top). At this time point, cells should have entered the S phase^36^. 87.6% of the differentiated synchronized cells re-entered the cell cycle during differentiation while 74% of the asynchronous control cells were in the S phase (Fig 4C, bottom) and were comparable to 2iLIF (Fig S4A). We also analysed the cell cycle distribution at 16 h after the shake-off and observed a higher enrichment of cells in G1 and G2M and less S phase compared to the asynchronous control in both differentiated cells and 2iLIF (Fig 4D and Fig S4B, respectively). Some of the G2M cells were potentially arrested in mitosis and did not re-enter the cell cycle after the nocodazole treatment. These G2M cells were generally bigger in nuclear area, we therefore measured the nuclear areas of cells that were EdU positive (i.e., re-entered the cell cycle) using EdU staining (data not shown). In the following experiments, the range of nuclear area of the EdU positive cells were used as a nuclear size range in nuclei segmentation of the smFISH data to exclude large, potentially arrested nuclei that did not re-enter the cell cycle after the shake-off.

**Fig 4:**
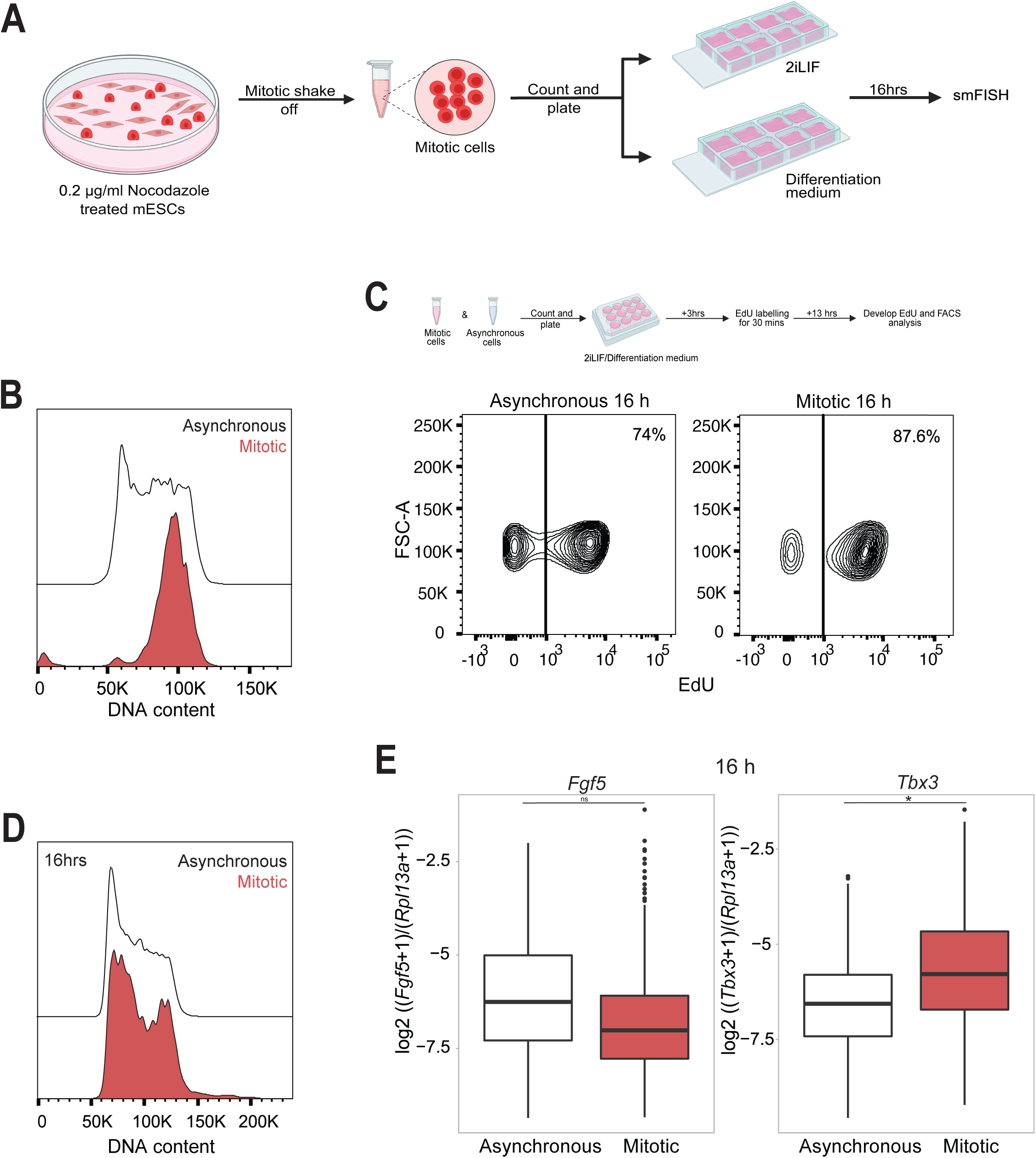
Cell cycle synchronization before induction of differentiation does not lead to a more synchronous exit from naive pluripotency. A. The experimental strategy of mitotic shake-off experiment followed by smFISH: mESCs were treated with 0.2 µg/ml Nocodazole to arrest cells in mitosis, 4 h later mitotic shake-off was performed. Collected mitotic cells were counted and plated in either 2iLIF or differentiation medium for 16 h B. Cell cycle distribution of synchronized mitotic cells after shake-off (mitotic), or DMSO treated (asynchronous) control cells, analyzed using Hoechst 33342 C. Incorporation of EdU measured at 16 h differentiation of asynchronous population (left) and cells released from mitosis (mitotic) (right). Schematic on top shows the experi­mental strategy: 30 mins EdU pulse was administered at 3 h post release from shake-off and 13 h later (total 16 h post release) EdU was developed and the percentage of positive cells was analyzed by FACS. The percentage of positive cells is indicated. D. Distribution of cell cycle at 16 h of differentiation after release from mitosis and asynchronous populations, analyzed using Hoechst 33342 E. *Rpl13a* normalized counts of *Fgf5* (left) and *Tbx3* (right) per cell for the asynchro­nous population and cells released from mitosis (mitotic) after 16 h of differentiation. calculated as log2 [(no. of *Fgf5* or *Tbx3* transcripts per cell +1) / (no. of *Rpl13a* tran­scripts per cell +1)]. F-test was performed on the log2 transformed values to test the significance of variability between asynchronous population and cells released from mitosis (mitotic). p<0.05 is considered significant. ns: non-significant. One experiment is shown (for the rest of the replicates see S6 I-L)

Next, we analysed the cell state at the single cell level using single molecule fluorescence in situ hybridization (smFISH) and quantified the number of selected transcripts per cell. The expression of three genes (*Fgf5* (formative marker), *Tbx3* (naive marker), and *Rpl13a (*housekeeping gene) were analysed. *Rpl13a* served here as an internal control for cell size as larger cells are known to produce more transcripts^37^. Indeed, the nuclear area correlated well with *Rpl13a* transcript levels per cell (Fig S5A, B) in both differentiated and 2iLIF asynchronous cells. The correlation was lower for the synchronized cells compared to the asynchronous cells because cells released from mitosis have a higher enrichment of smaller cells/nuclei (G1) (Fig S5C,D). However, no correlation of nuclear size to *Fgf5 (*Fig S5E,F) or *Tbx3* (Fig S5I-J) counts per cell was observed in the asynchronous differentiated or 2iLIF population. Similarly, we did not observe a correlation between nuclear size and *Fgf5* or *Tbx3* expression for cells synchronized to mitosis either in differentiation or under 2iLIF conditions (*Fgf5:* Fig S5G,H, *Tbx3*: Fig S5K,L).

The homogeneity of the cell state transition was assessed by comparing the variance in the expression of cell state markers across a population of synchronized or unsynchronised cells that were induced to differentiate. The F-test is used to statistically compare variances but it assumes normality of the data. Therefore, we used log2 transformed *Rpl13a* normalized counts of *Fgf5* and *Tbx3* that we tested for normality using a QQ-plot (*Fgf5*: Fig S6A-D, *Tbx3*: Fig S6E-H). For the differentiated cells, there was no significant difference in the variance of the normalized *Fgf5* counts per cell in the synchronized compared to asynchronous control in most of the replicates (Fig 4E, Fig S6I, (left panel)). One of our replicates of the cells released from mitosis had significantly lower variance compared to asynchronous control for *Fgf5* (Fig S6K (left)). For *Tbx3* most of the replicates had no significant difference in variance (S6 I,K (right)), and one replicate had a significantly higher variance of expression (Fig 4E (right)) in mitotic vs. asynchronous populations. In the undifferentiated cells the variance of the normalized *Fgf5* was not significantly different in the cells released from mitosis compared to the asynchronous population in all the replicates (Fig S4C(left), S6 J,L (left)), while for *Rpl13a* normalized counts of *Tbx3*, most replicates have no significant difference in variation in synchronized compared to asynchronous (Fig S4C (right), Fig S6J (right)), except for one replicate which was significantly higher in variation in the mitotic compared to asynchronous (Fig S6L (right)).

Taken together these results show that the expression of *Fgf5* and *Tbx3* across the population of differentiating cells is not consistently more homogeneous in the mitotic release population compared to the asynchronous population.

### Differentiation delay phenotypes do not show any cell cycle bias

We reasoned that if the cell cycle phase plays a strong role during the exit from naive pluripotency, then an extension in the cell cycle length should lead to a delay in differentiation. Boileau *et al.* recently investigated the differentiation behaviour of MLL3/4 DKO and MLL3 KO mESCs^38^. The KOs showed a strong delay in proliferation, however, loss of the chromatin modifiers had little effect on the overall exit from naive pluripotency, suggesting that extension of cell cycle length does not interfere with differentiation. We, therefore, asked whether mESCs with a differentiation delay showed a different cell cycle profile. We analysed *Tcf7l1* and *Rbpj* knockout cell lines (*Tcf7l1^-/-^* and *Rbpj^-/-^*) in a Rex1-GFPd2 reporter cell line background (called RC9; used as the WT control) that were previously characterized with a strong differentiation delay^12^. First, we confirmed the differentiation delay in our differentiation assay by inducing differentiation of the *Tcf7l1*^-/-^ and *Rbpj^-/-^,* and RC9 cells as control and measuring GFPd2 expression by FACS. We collected the cells at 6 h, 12 h, and 48 h. The WT showed a clear reduction in GFPd2 positive cells at 48 h while the *Rbpj^-/-^*and *Tcf7l1*^-/-^ were still enriched for GFPd2 positive cells, indicating a delay in differentiation (Fig S7 A, B, C (WT), D, E, F (*Rbpj^-/-^*), G, H, I (*Tcf7l1^-/-^*)). This was corroborated by analysing MULTI-seq data from the differentiation of *Tcf7l1*^-/-^ and *Rbpj^-/-^,* compared to WT cells (Fig S8 A (6 h), B (12 h), C (48 h)) by PCA. We then analysed the cell cycle profile of the cells using Hoechst 33342 staining of cells at the indicated time points in differentiation. Compared to the WT control, both knockout cell lines have very comparable cell cycle profiles at 6 h, 12 h, and 48 h indicating no cell cycle bias for slower differentiating cells (Fig 5A, C, E). Classification of differentiating cells at 6 h, 12 h, and 48 h into the different cell cycle phases using Cyclone showed no differences in pseudotime for *Tcf7l1*^-/-^ and *Rbpj^-/-^,* compared to WT cells (Fig 5B, D, F). Taken together our data shows that slower differentiation cannot be attributed to cell cycle differences.

**Fig 5:**
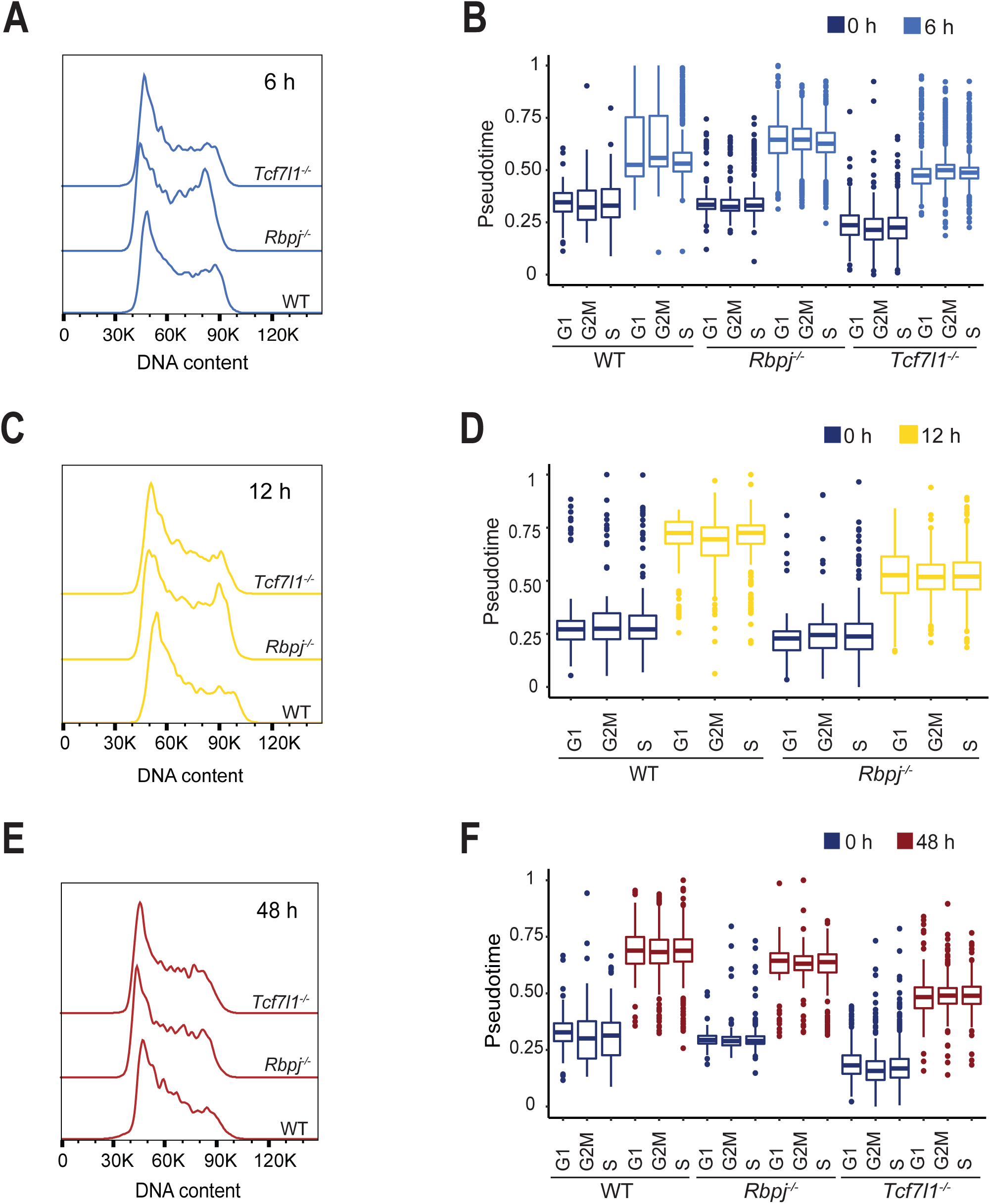
Differentiation delay mutants do not show any cell cycle differences compared to WT upon exit from naive pluripotency. A. Cell cycle distribution of *Tcf7l1^-/-^, Rbpj^-/-^* and WT (Rex1-GFPd2) differentiated for 6 h measured by Hoechst 33342 staining B. Distribution of pseudotime of WT (Rex1-GFPd2), *Rbpj^-/-^, and Tcf7l1^-/-^* classified into G1, S and G2M, using Cyclone (Scialdone et al. 2015) cultured in 2iLIF (0 h) and differentia­tion medium for 6 h C. Cell cycle distribution of *Tcf7l1^-/-^, Rbpj^-/-^* and WT (Rex1-GFPd2) differentiated for 12 h measured by Hoechst 33342 staining D. Distribution of pseudotime of WT (Rex1-GFPd2) and *Rbpj^-/-^* classified into G1, S, and G2M using Cyclone cultured in 2iLIF (0 h) and differentiation medium for 12 h. *Tcf7l1^-/-^* not shown because of loss of cells during cell pooling for 10X Genomics scRNA-seq E. Cell cycle distribution of *Tcf7l1^-/-,^ Rbpj^-/-^* and WT (Rex1-GFPd2) differentiated for 48 h measured by Hoechst 33342 staining F. Distribution of pseudotime of WT (Rex1-GFPd2), *Rbpj^-/-^* and *Tcf7l1^-/-^* classified into G1, S and G2M using Cyclone cultured in 2iLIF (0 h) and differentiation medium for 48 h

## DISCUSSION

In this study, we demonstrate that the exit from naive pluripotency is a continuous cell state transition with no distinct intermediate cell states. Differentiation, however, occurs asynchronously. Though several studies point to the G1 phase as a window of opportunity to initiate cell state changes in pluripotent cells, it remained unclear whether it dictates the timing of exit from naive pluripotency in mESCs ^17,26–28,33,36^. The majority of previous studies were performed in hESCs. Here, the cell cycle state determines cell fate propensity through G1 specific activation of TGF-beta signalling via Cyclin D^26^. However, hESCs exist in the primed state of pluripotency while mESCs cultured in 2iLIF exist in the naive state of pluripotency^2,39^. The naive state resembles the pre-implantation epiblast cells while the primed state represents the late post-implantation epiblast^11^. In mice, the primed state is captured in epiblast stem cells (EpiSCs) that are developmentally distinct from naive cells: the latter can contribute to chimeras when injected into the blastocyst while the former cannot^40^. Interestingly, mESCs exhibit a distinct cell cycle regulation compared to hESCs. hESCs show cell cycle dependent fluctuation of CDK2^24,41^ and express cyclin Ds (D1, D2 and D3) during late G1-S, while mESCs exhibit phase specificity only for CyclinB/Cdk1^42^. Cyclin D (Ccnd1 and Ccnd2) is expressed in naive mESCs throughout the cell cycle with a slightly elevated expression at late G1^43^. These differences between the human and mouse ESCs indicate that there could be a different relation between the cell cycle and cell state in mESCs. In support of this, it has been suggested that G2M diverts mESCs grown under serum/LIF conditions into naive and formative states of pluripotency due to heterogeneity in the *Esrrb* levels (*Esrrb* high and low states)^44^. Retinoic acid driven extra embryonic endoderm (XEN) differentiation of mESCs has also been linked to G2M specific expression of Esrrb^45^.

To address if the cell cycle is a major source of asynchrony in the exit from naive pluripotency, we first performed a time course scRNA-seq experiment to capture the transition. The exit from naive pluripotency is a continuous cell state transition, however, it is asynchronous in nature. Further differentiated cells at any experimental time point do not belong to any particular cell cycle phase, indicating that the cell cycle and cell state are uncoupled during the exit from naive pluripotency. Since scRNA-seq data does not provide information on cell cycle phases at the onset of differentiation, we resorted to cell cycle synchronization experiments. At 12 h and 16 h of differentiation, the formative and naive pluripotency marker expression levels were comparable to the asynchronous control (see also Waisman *et al*. ^36^). Direct comparison of the expression of pluripotency and differentiation markers to the asynchronous differentiated control revealed that most genes do not show differences in expression across the different cell cycle sorted populations. Particularly, *Fgf5* expression was cell cycle regulated at both 12 h and 16 h (see also Waisman *et al* ^36^). Furthermore, *Otx2* showed significantly lower expression than the asynchronous control at 12 h and 16 h in a cell cycle-dependent manner indicating that it might be a cell cycle regulated gene. OTX2 has been shown to regulate G1-S transition in medulloblastoma^46^ and is expressed in the G2M phase of the last cell cycle during mouse retinal development^47^, however, it has not been implicated to have a cell cycle dependency in the exit from naive pluripotency. Next, we showed that starting differentiation from a synchronized population of mESCs does not lead to increased homogeneity in the exit from naive pluripotency (Fig 4E). *Tbx3* expression at 16 h is significantly more variable in the population released from mitosis compared to the asynchronous population potentially because it is a lowly expressed gene during differentiation; consistent with reports of lowly expressed genes being more heterogeneous in expression^48,49^.

Asynchronous exit from naive pluripotency has been attributed to cell variability in ERK signalling dynamics via Nanog expression control^50^. On the contrary, dynamics (sustained vs. transient activation) in ERK signalling was reported to have surprisingly little correlation to the exit from naive pluripotency, also measured by Nanog levels. Instead, local cell density influenced the dynamics: cells with many neighbours had less ERK activity, albeit to similar levels across the cells^51^. ERK signalling dynamics itself have been reported to be cell cycle dependent with sustained activity at the beginning of the cell cycle compared to later phases of the cell cycle in mESCs cultured in 2i, however with little dependency on Nanog states (low vs high) ^52^. Nevertheless, we do not observe a link between the cell cycle and the timing of exit from naive pluripotency, therefore, cell cycle regulated ERK activity does not seem to influence the decision to differentiate for mESCs cultured in 2iLIF.

Thymidine glycosylase (TDG) is a cell cycle dependent enzyme enriched in G1 phase that causes transcriptional heterogeneity in mouse embryonic stem cells by acting as a co-factor to p53 target genes and facilitates cardiac fate specification when induced to form embryoid bodies (EB) from G2M phase (TDG less) compared to G1^53^. Nevertheless, the effect on cell fate for heterogeneous TDG populations was observed after 11 days of EB differentiation, limiting our understanding of the influence of the cell cycle during the onset of differentiation.

Our study shows the lack of involvement of the cell cycle in the asynchrony during the exit from naive pluripotency and thereby supports recent work by Strawbridge S. *et al*.^17^ Here, variable lag periods in the downregulation of a pluripotency marker Rex1 were observed, irrespective of the cell cycle phase. Evidence for the lack of cell cycle control in cell fate decisions also exists in other model organisms. Blocking the cell cycle at S phase or in G2M phases did not affect cell type specification during zebrafish development between early gastrulation to somitogenesis^54^. In *C. elegans* embryo gut specification, the transcriptional onset of genes was shown to be uncoupled from cell division timing^55^.

This opens doors to explore other mechanisms like differential signalling responses, local cellular environment, and transcriptional bursting as underlying reasons for asynchrony in the exit from naive pluripotency. The local environment in the form of differential localization of signalling ligands in the culture medium could lead to different downstream responses. mESCs exposed to localized Wnt signalling divided asymmetrically where proximal cells adopted a naive ESC fate while the distal cells expressed EpiSCs markers ^51^. Sibling cells show synchronous dynamics of exit from naive pluripotency^17,33,52^, indicating that neighbouring cells have similar transcriptional profiles and thereby influence heterogeneity. The sources of cellular variability have been ascribed to transcriptional bursting, since transcription is discontinuous with periods of active transcription interspersed by periods of inactivity^56^. Transcriptional bursting regulates phenotypic switching in eukaryotic cells^57^, mouse erythropoiesis^58^, and response to estrogen in breast cancer cells^59^. Therefore, we speculate that transcriptional heterogeneity that arises due to differential bursting between individual mESCs could render some cells more pluripotent than others and thereby lead to different responses to differentiation cues.

## Supporting information

supplement table primer

## Acknowledgments

We would like to thank all members of the Buecker lab for discussions and feedback throughout the project, Martin Leeb, and the members of his lab for continuous support and critical feedback in shared group meetings. In addition, we would like to thank Julia Naas for the discussions and critical feedback on statistical analysis. The BioOptics-FACS and BioOptics-Lightmicroscopy facilities at the Max Perutz labs were instrumental in the success of this project. Some figures were created with BioRender.

scRNA-seq and Quant-seq was performed by the Next Generation Sequencing facility at the Vienna BioCenter Core Facilities (VBCF), a member of the Vienna BioCenter (VBC), Austria.

The work was supported by the Austrian Science Fund FWF (P34123 to CB and W1261 DK SMICH) and Uni:Docs fellowship from the University of Vienna to M.R.

## METHODS

### mESC maintenance

For regular maintenance mouse embryonic stem cells (R1 and the other cell lines where applicable) were cultured in base medium HyClone DMEM/F12 without Hepes (Cytiva), with 4 mg/mL AlbuMAX™ II Lipid-Rich Bovine Serum Albumin (GIBCO™), 1× MACS NeuroBrew-21 with Vitamin A (Miltenyi Biotec), 1× MEM NEAA (GIBCO™), 50 U/mL Penicillin–Streptomycin (GIBCO™), 1 mM Sodium Pyruvate (GIBCO™), and 1× 2-Mercaptoethanol (GIBCO™), supplemented with 3.3 mM CHIR-99021 (Selleckchem), 0.8 mM PD0325901 (Selleckchem), and 10 ng/mL hLIF (provided by the VBCF Protein Technologies Facility, https://www.viennabiocenter.org/facilities/) to complete the 2iLIF self renewal medium. Cells were cultured on Greiner Bio-One CELLSTAR Polystyrene 6-well Cell Culture Multiwell Plates coated first with Poly-L-ornithine hydrobromide (6 μg/mL in 1xPBS (Sigma-Aldrich), 1 h at 37℃, Sigma-Aldrich) followed by Laminin from Engelbreth-Holm-Swarm murine sarcoma basement membrane (1.2 mg/mL in 1xPBS, 1 h at 37℃, Sigma-Aldrich). Cells were routinely passaged in a 1:6 ratio every 2 days using 1× Trypsin–EDTA solution (Sigma) at 37°C and the reaction was stopped using 10% fetal bovine serum (FBS, Sigma) in base medium.

### Hoechst 33342 staining and cell cycle phase sorting

10 million mESCs (R1) were seeded onto two, 15 cm dishes (CytoOne 150 x 20 mm Tissue Culture Dish) each 24 h before collection for 10 mg/mL Hoechst 33342 (Thermo Fisher Scientific) staining. Cells were washed twice with PBS and trypsinized for 10 mins at 37℃. Trypsinization was stopped using 10% FBS in base medium, cells were counted and resuspended in 15 μg/mL Hoechst 33342 solution (in 10% FBS in base) at 37℃ for 20 mins at a concentration of 1000 cells/ul. Next, the cells were centrifuged and resuspended in 15 μg/mL Hoechst 33342 solution at a concentration of 41,000 cells/ul, passed to a round bottom Falcon 5mL tube through with a cell strainer cap (Corning), and sorted using BD ARIA cell sorter using the UV filter. Initially, 2-3 million cells were sorted without gates (asynchronous control) followed by sorting 2 million cells gated based on their DNA content into G1, S, and G2M phases. The cells were counted again and 100,000 cells were seeded onto 5 μg/mL Humanplasma Fibronectin (Sigma Aldrich,1 mg/mL) in PBS coated (1 hour at 37℃) 12 well plates (Greiner Bio-one, Cell culture multiwell plate, 12 well, PS) either in differentiation medium (base medium supplemented with 12 mg/mL Recombinant Human FGF-basic (PEPROTECH) and KnockOutTM Serum Replacement (1:100, GIBCO™), hereafter referred to as FK medium) or 2iLIF as undifferentiated control.

### Propidium Iodide (PI) staining

PI staining was performed to analyse cell cycle purity at 12 and 16 h after differentiation (and 2iLIF control) of the cell cycle sorted populations. Cells left over after seeding (∼10^5^ cells) were pelleted down at 1000 rpm for 2 mins and resuspended in 1xPBS and 100% ice-cold ethanol was added dropwise while vortexing such that the final concentration of ethanol was 70%. The samples were incubated for at least 30 mins or overnight until PI staining was performed. Then, they were centrifuged for 5 mins at 500 x g and the supernatant was discarded. Pellets were resuspended in 50 μL of 100 μg/mL RNAse A (Thermo Scientific, 10 mg/mL) and incubated for 5 mins at RT. To this 250 μL of 50 μg/mL propidium iodide (Sigma Aldrich, 10 mg/mL) was added and incubated for 5 mins in the dark, transferred through a cell strainer cap (Corning) and analyzed at the BD ARIA cell sorter. Data was analysed using FlowJo 10.8.1 (see cell cycle measurement using Hoechst 33342 for details of analysis).

### RNA extraction and qPCR

Cells were lysed after 12 h or 16 h using 500 μL pepGOLD TriFast ™ reagent (VWR) and stored at -80 until the RNA extraction process was resumed. RNA extraction was performed according to the manufacturer’s protocol by phenol chloroform extraction followed by isopropanol precipitation, and 75% ethanol washes. 500ng of RNA was used for cDNA synthesis using SensiFAST ™ cDNA Synthesis kit (LabConsulting) according to the manufacturer’s protocol. The cDNA was diluted 1:2 in nuclease free water (SIGMA-Aldrich) and real time quantitative PCR (RT-qPCR) was performed using SensiFAST™ SYBR No-ROX kit (LabConsulting). 0.5 μL of cDNA was combined in a 10 μL reaction with 125 nM RT-qPCR primers designed using Primer3^60^. qPCR was performed in 3 technical replicates and an average Cq value of individual primers in each population (G1, S, G2M, asynchronous) was calculated. Delta-Delta Ct method was followed for further analysis. First, the average Cq value from each gene was subtracted from the average Cq of *Rpl13a* (housekeeping gene) (called ΔCT). Then normalization was performed for each gene in two different ways: 1) to the undifferentiated counterparts of each population, and 2) to the asynchronous differentiated population. In both cases, ΔCT values of each gene within a population were subtracted from the corresponding ΔCT of the 1) undifferentiated of each population or 2) asynchronous differentiated population. This value is called the ΔΔCT. Finally, to get the fold change of expression levels of each gene 2^ΔΔCT was calculated. For statistical analysis, we performed the unpaired Wilcoxon test in R Studio Version 2022.12.0+353 (R 3.6.2) on log2 [2^(ΔΔCT)] values calculated. A p-value less than 0.05 was considered significant. The box plots were generated using the ggplot2 (version 3.3.5) package in R studio.

### Mitotic shake-off

The protocol was adapted from Effie Apostolou’s lab^29^. 7.5 million R1 mESCs were seeded onto two, 15 cm dishes (CytoOne 150 x 20 mm Tissue Culture Dish) in 2iLIF. 24 h later the cells were washed twice with PBS and replaced with 20 mL of either 0.2 μg/mL Nocodazole in 2iLIF (SIGMA 1mg/mL stock) or DMSO in 2iLIF. 4 hours later the Nocodazole containing plate was washed to detach the rounded mitotic cells gently using a 10 mL pipette. The plate was washed once again with 20 mL PBS to collect any mitotic cells left behind. The DMSO containing plate was trypsinized for 10 mins using 10 mL trypsin and the reaction was stopped using 10 mL 10% FBS in base medium. To maintain consistency, 20 mL PBS was used to wash the DMSO plate to obtain any cells left behind. The samples were centrifuged for 5 mins at 1000 rpm (this was maintained in all further centrifugation steps), resuspended in 1mL PBS, and counted. From the DMSO sample, the same number of cells as in the mitotic sample were used for the experiment as there were excess cells. Then, to prevent cell clumping due to extruded DNA from broken cells, they were incubated in 100 μg/mL DNAse 1 (1 mg/mL, Merck) in a concentration of 1000 cells/μL for 15 mins at RT. Then, the samples were washed 3 times in ice-cold 0.5% Bovine Serum Albumin (BSA) (Sigma-Aldrich) in PBS (to prevent cell clumping) by centrifugation for 5 mins at 1000 rpm in between washes. 100 μg/mL DNAse1 incubation step was repeated to clear any further cell clumping due to extruded DNA due to cell damage during centrifugation. Finally, the two populations; DMSO (asynchronous control) and the Nocodazole-treated mitotic cells were counted again and 60,000 cells were plated onto 5 μg/mL Humanplasma Fibronectin (Sigma Aldrich,1mg/mL) in PBS coated (1 hour at 37℃) 8 chambered cover glass (µ-Slide 8 Well Glass Bottom, IBIDI) (200 μL added per well). Each sample was plated in FK medium and 2iLIF medium, fixed 16 h later for smFISH. One well each of the DMSO treated sample was seeded in FK medium and 2iLIF to use as a “no probe” negative control in the smFISH and processed along with the other samples.

### Cell cycle measurement using Hoechst 33342

To test the purity of obtained mitotic cells and DMSO control (asynchronous) cells, they were counted and stained with 15 μg/mL Hoechst 33342 solution as described above. After staining, cells were centrifuged for 2 mins at 1000 rpm and resuspended in 300 μL of 15 μg/mL Hoechst 33342 solution, strained through the cell strainer cap (Corning), and fluorescence was measured in the BD ARIA cell sorter using the UV filter. The cell cycle analysis was done by first performing doublet discrimination. The first gate was set using FSC-A vs. SSC-A followed by a second more stringent gate on this population using SSC-A vs. SSC-H. These were identified as single cells and the cell cycle profile was plotted as a histogram of the DAPI channel that measured the Hoechst 33342 dye. All FACS analyses were carried out using FlowJo 10.8.1.

To measure the cell cycle phase at 16 h post release from mitotic shake-off, 210,000 cells each of asynchronous and mitotic cells were seeded (same density as 60,000 cells per well of the 8 chambered cover glass) onto a 5 μg/mL Humanplasma Fibronectin (Sigma Aldrich,1mg/mL) coated in PBS (1 hour at 37℃) (500 μL added per well) 12 well plates (Greiner Bio-one, Cell culture multiwell plate, 12 well, PS). At 16 h post-release, the asynchronous population and cells released from mitosis were trypsinized, counted, and stained with 15 μg/mL Hoechst 33342 solution as described above. After staining cells were centrifuged for 2 mins at 1000 rpm, resuspended in 300 μL Hoechst 33342 solution, and analyzed using the BD ARIA cell sorter using the UV filter.

Cell cycle measurements of *Tcf7l1^-/-^*, *Rbpj^-/-^ (*kindly provided by the lab of Martin Leeb), and WT (Rex1-GFPd2) cells were also performed as described above after inducing differentiation. 70,000 cells of each cell line were seeded on 5 μg/mL Humanplasma Fibronectin (Sigma Aldrich,1mg/mL) in PBS coated (1 hour at 37℃) (500 μL added per well) 12 well plates (Greiner Bio-one, Cell culture multiwell plate, 12 well, PS). 16 h later cells were washed twice with PBS and differentiation was initiated as described above. After 6, 12, and 48 h cells were trypsinized and Heochst 33342 stained and analysed by FACS as described above. The undifferentiated samples were also stained with Hoechst 33342 and fluorescence was measured using FACS. The cell cycle profile was analyzed using FlowJo 10.8.1.

### Single molecule fluorescence in situ hybridization (smFISH)

To perform smFISH after mitotic shake-off, mitotic released and asynchronous control cells were fixed in 4% paraformaldehyde (16% paraformaldehyde solution, MP Biomedicals) for 45 mins at room temperature. Cells were then washed twice in PBS and stored in the fridge until the experiment proceeded. The QuantiGeneⓇ ViewRNA™ ISH Cell Assay kit was used according to the manufacturer’s protocol. Fixed samples were washed twice with PBS and permeabilized using the company provided Detergent Solution QC for 5 mins at room temperature followed by two PBS washes. Hybridization was performed using the following probes after diluting them 1:100 in Probe Set Diluent QF: *Fgf5* mRNA (Cy3) (View RNA *Fgf5* Type 1 (mRNA): VB1-18683 LOT 346763-000), *Tbx3* (GFP) (View RNA *Tbx3* Type 4: VB4-18685 LOT 268746-000), *Rpl13a* (Cy5) (View RNA MO *Rpl13a* Type 6: VB6-15315 LOT 117193179) for 3 h at 40℃. This was followed by hybridization using the PreAmplifier mix (PreAmplifier Mix was diluted in pre-warmed Amplifier diluent QF in the ratio 1:25) for 30 mins followed by three wash buffer washes (made as directed by the company using provided component Wash Buffer components 1 and 2) for 2 mins each. Similarly, Amplifier Mix, and Label Probe Mix were hybridized. The cells were then washed with the wash buffer three times and left in the last wash for 10 mins. Finally, DAPI staining was performed using 1x DAPI in PBS diluted from 100x DAPI provided by the company. Cells were incubated in DAPI for 1min washed twice in PBS and stored at 4°C in 1:50 Penicillin-Streptomycin (Thermo Fischer Scientific 5000U/mL) and Gentamicin solution (Sigma Aldrich, 10mg/mL solution) in PBS until imaged. Prior to imaging, 300 μL antifade solution (Glox buffer from Raj Lab protocols) was added to the samples. Imaging was performed on Zeiss Cell Discoverer 7 with Plan-Apochromat 50x/1.2W autocorr, WD 0.84mm objective. For each sample, 20-30 positions were imaged as a stack containing 27 slices spaced 0.5um apart.

### Image analysis

Images were maximum intensity projected using ImageJ (Version 2.1.0) and channels were split using a publicly available Macro script. Images were further analyzed using Cell Profiler (version 4.2.1) using a custom-made pipeline. For background subtraction from channels, a Gaussian filter (size 120, for nuclei channel and 20 for channels detecting *Fgf5, Rpl13a* and *Tbx3*) was applied to the image using the Gaussian filter module, and the filtered image was subtracted from the raw image. To detect the mRNA spots in each channel, a threshold intensity was determined from the background subtracted negative control (no-probe) images. The spots in the negative control images were detected using a manual threshold in the Identify PrimaryObjects module and their intensities were quantified. The 80^th^ percentile of the median intensity values was used as a threshold to detect spots (using the Identify PrimaryObjects module) in the mitotic and asynchronous control images (differentiated and undifferentiated). To associate transcripts to a cell, first, nuclear segmentation was performed using the IdentifyPrimaryObjects module, and the nuclei were used as a seed to expand an area around the nuclei to mark the cell boundary, using the IdentifySecondaryObjects module (Distance N method: 40 pixels). Using the RelateObjects module, transcripts found within the area of a cell were associated with it. The nuclei segmentations were manually inspected and clumped / half nuclei detections were later removed from the data.

### Data analysis of smFISH data

To plot the number of transcripts per cell it was first normalized to *Rpl13a*. To do that, a pseudocount of 1 was added to both *Fgf5*, *Tbx3,* and *Rpl13a* since some of these values were 0. Then, the corresponding gene counts were divided by the newly calculated *Rpl13a* values within each cell.

The normalized values were log2 transformed and plotted for the asynchronous and mitotic released population. F- test was performed on the log2 transformed values using R Studio Version 2022.12.0+353 (R 3.6.2) using ggpubr (version 0.4.0) and ggplot2 (version 3.3.5). Since F-test assumes normality of the data, we tested it using a QQ - plot that was plotted using the log2 transformed values in R Studio using the quantile-quantile plot commands (qqnorm and qqline). The nuclear area was also quantified using the MeasureObjectSizeShape module. The “shape area”, which is the sum of the pixels within the segmented nuclear region, was quantified and plotted against the number of transcripts of *Fgf5*, *Tbx3,* and *Rpl13a* per cell using the ggplot2 and ggpubr packages mentioned above.

### EdU staining

EdU Assay/EdU Staining Proliferation Kit (iFluor 647) from Abcam (ab222421) was used to perform the assay. First, mitotic shake-off was performed, and mitotic and asynchronous cells (210,000) were seeded in FK medium and 2iLIF as described above onto 5 μg/mL Humanplasma Fibronectin (Sigma Aldrich,1mg/mL) in PBS coated (1 hour at 37℃) (500 μL added per well) 12 well plates (Greiner Bio-one, Cell culture multiwell plate, 12 well, PS), each with a well of cells for no EdU control as well. 3 h post release of the cells, the cells were incubated for 30 mins with 10μM EdU made in 2iLIF / FK medium followed by washing it off two times using PBS and replacing it with respective media. 13 h later, cells were trypsinized and trypsin was stopped using 10% FBS in base. All the instructions from the kit were followed after the collection of cells. Samples were resuspended in base medium and washed two times in the recommended wash buffer (3%BSA in PBS) by centrifuging for 5 mins at 300 x g at 4℃ (this was followed in all subsequent centrifugation steps). 4% paraformaldehyde (from 16% paraformaldehyde, MP Biomedicals) was used to fix the cells in suspension by incubating at room temperature for 20 mins. Fixative was removed followed by two wash buffer washes. Permeabilization was performed for 30 mins at room temperature using 1X Permeabilization buffer made from 10X Permeabilization buffer provided by the company followed by EdU development. 1mL of EdU reaction mix was made according to the company instructions and reagents and then samples were incubated in 100 μL of reaction mix for 30 mins in the dark. Samples were centrifuged and washed twice in 1x Permeabilization buffer and resuspended in 300 μL PBS. Samples were passed through a 5 mL tube with a cell strainer cap (Corning) and FACS fluorescence was measured using the VIOLET filter setting of the BD ARIA cell sorter. FACS data was analysed in FlowJo 10.8.1 as described, for Hoechst 33342 but data was represented using a PE-A vs. FSC-A plot.

### 10X Genomics scRNA-seq with MULTI-Seq

Cells were harvested by trypsinization as described above. Cell pellets were resuspended in PBS to remove residual FCS and counted with a CASY cell counter (Biovendis). Labeling of cells with MULTI-Seq barcodes was performed as described^34^. In short, 0.5 million cells per sample were resuspended in PBS and incubated with barcode + lipid anchor mix for 5 mins of ice. Co-anchors were added for an additional 5 mins, then the reaction was quenched with 1% BSA/PBS. Cells were washed two times in 1% BSA/PBS and finally resuspended in 0.04% BSA. All samples were combined, filtered through FACS strainer cap tubes, and adjusted to 1 million cells/mL. Only pools with > 80% cell viability were used for subsequential 10X library preparation according to the manufacturer’s instructions. Libraries for MULTI-Seq identifications were prepared alongside the cDNA libraries sequenced on Illumina NextSeq550 or NovaSeq For validation of the specificity of MULTI-Seq, ATTO-488 and ATTO-590 conjugated barcodes were used, as described (McGinnis *et al.* 2019). Cells were labelled with barcodes as described above, but labelling was validated by FACS.

### Data Analysis scRNA-seq

cDNA reads were mapped to *mm10* and demultiplexed into droplets using *Cellranger*. MULTI-Seq barcodes were mapped to droplets and counted using *CITE-Seq-Count* and the classification of cells was performed following the Seurat vignette “Demultiplexing with Hashtag oligos”. Basic quality control on library sizes and mitochondrial read content was performed with *scater.* For normalization and log conversion of gene expression counts *scran* was used. HVGs were calculated by modelling mean-variance relationships per gene and selecting the top 500 genes. We then based dimension reduction on these selected genes

We used *cyclone*^35^ to infer cell cycle phases. To regress for cell cycle, we used the determined cell cycle phase as block while modelling mean-variance relationships and selected the resulting top 500 HVGs. For pseudotime analysis, we used *slingsho*t based on the indicated PCA dimension reductions. We scaled and reverted the resulting pseudotime values so that 0 corresponds to the start and 1 to the end point of differentiation.

We used *ggplot2* to make boxplots and all other data visualizations. Boxplots show the mean as central line, 25^th^ and 75^th^ percentiles as hinges, and 1.5* inter-quartile range as whiskers.

### Bulk RNA seq

RNA QuantSeq was prepared according to the manufacturer’s instructions (Lexogen 3’ mRNA Seq Library Prep Kit). 500 ng RNA was used as the starting material. qPCRs were used to evenly multiplex samples and final libraries were quantified by NEBNext Library Quant Kit for Ilumina (#E7630S). QuantSeq RNA seq data was processed following Lexogen’s standard pipeline on bluebee. Adapter contamination was removed with bbmap, mapped using STAR and counted with HTScount. Downstream analysis was performed in R using DESeq2 (1.30.1) and pheatmap (1.0.12). We used DESeq2 to calculate differential gene expression between ESCs and EpiLCs and selected based on the adjusted p-value the top 500 most differentially expressed genes.

---

**Fig S1 (related to Fig 1):**
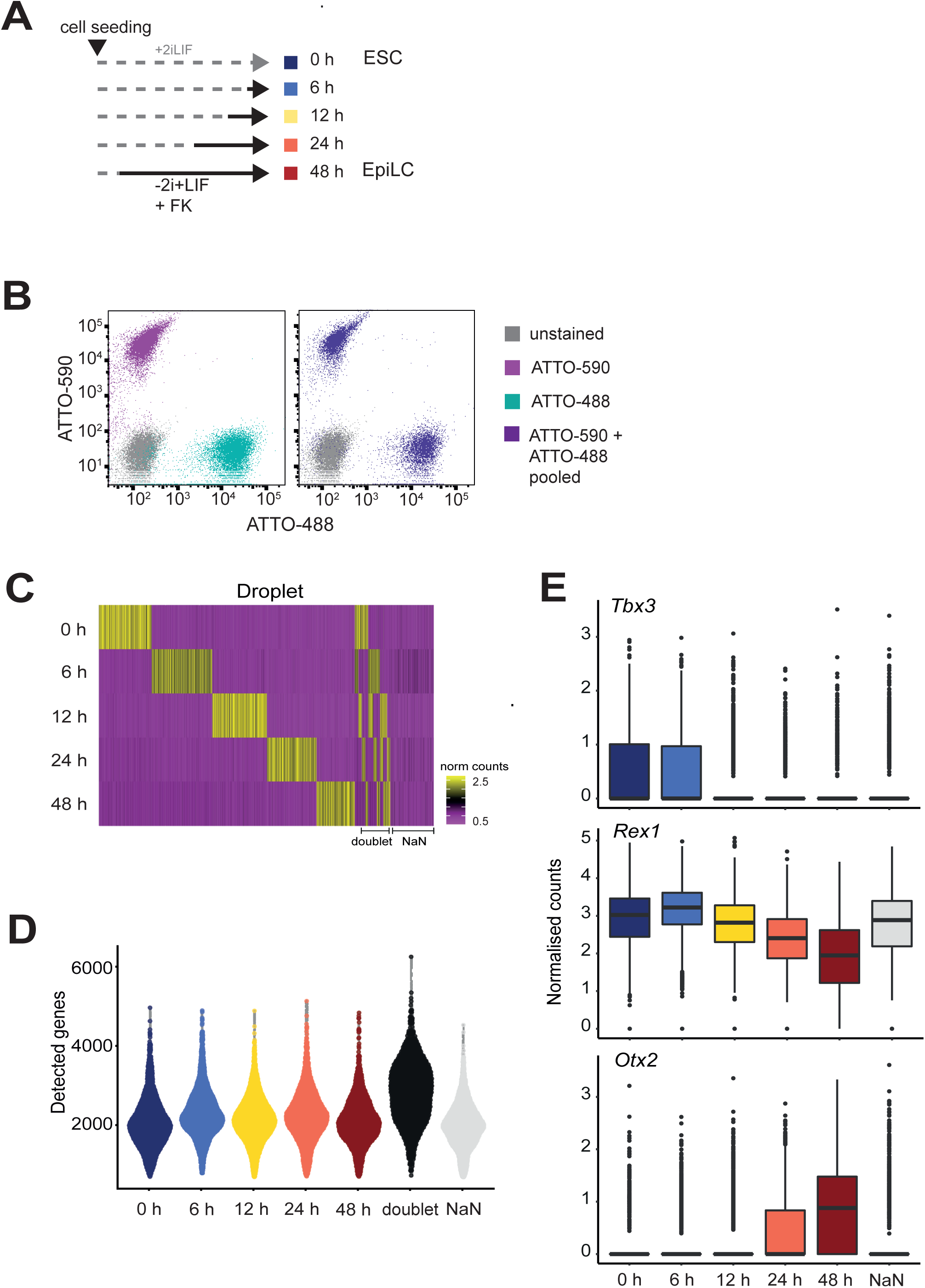
A. Experimental setup of scRNA-seq experiment time course: Cells were seeded at the same time, differentiation was initiated at the indicated time points (undifferentiated control was maintained in 2iLIF), and all samples were collected at the same endpoint B. Validation of specificity of MULTI-seq barcodes using fluorescently labeled DNA barcodes via FACS; left: individual staining with two different fluorescent barcodes; right: cells were pooled after staining with two different fluorescent barcodes C. Identification of samples by MULTI-seq: Columns represent filtered droplets and rows represent normalized counts for different barcodes, grouped based on the time point of collection. Most droplets are associated with a single barcode while some have more than one barcode associated and are therefore labelled as doublets. Those associ­ated with no barcodes are labeled as NaN. D. Number of genes detected per cell before quality-based filtering of cells at the indicated time points E. Normalized counts per cell of *Tbx3, Rex1*, and *Otx2* at the indicated time points

**Fig S2 (related to Fig 1):**
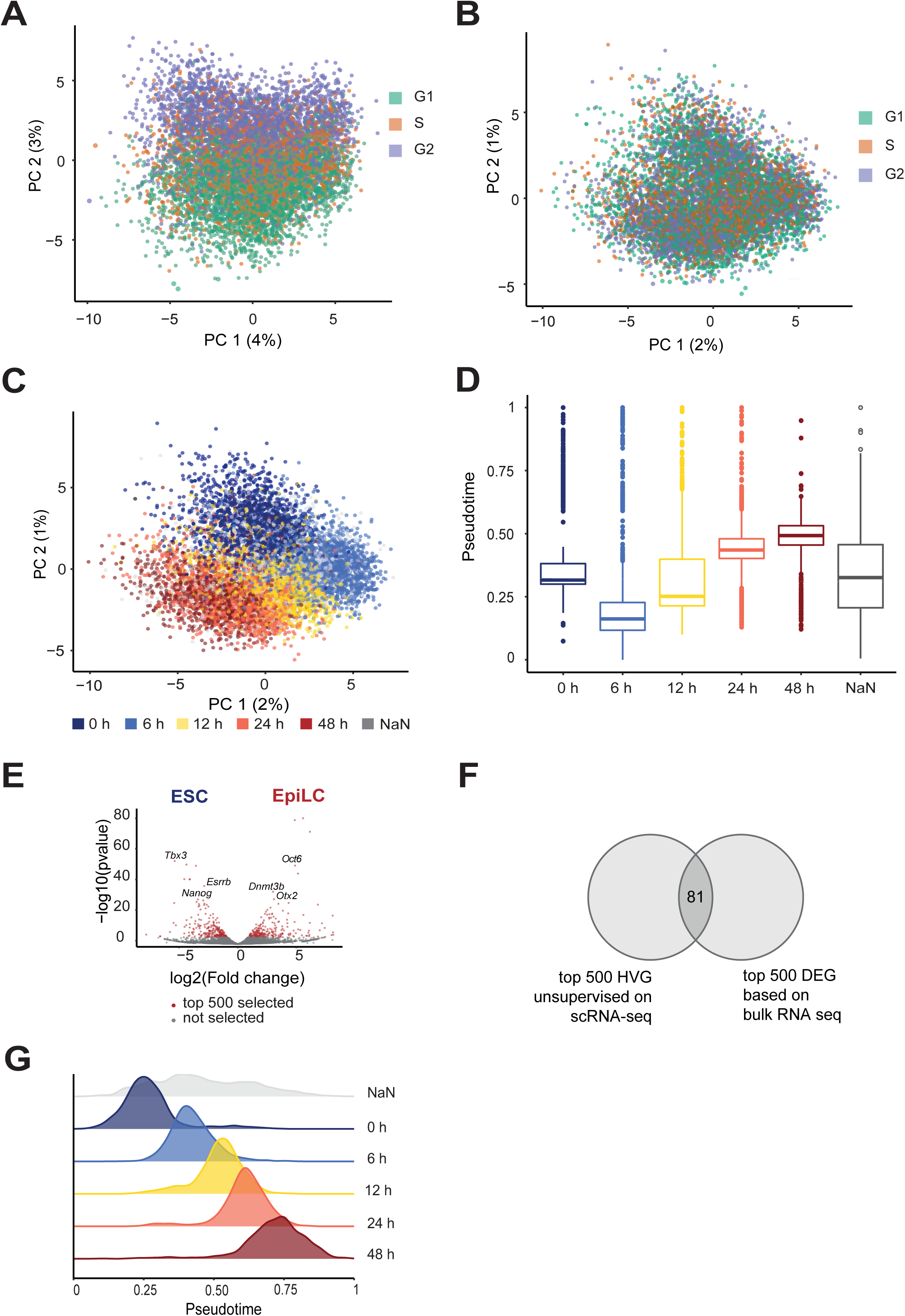
A. PCA of scRNA-seq data (as in Fig 1A) overlaid with cell cycle phases detected using Cyclone (Scialdone et al. 2015) as indicated by the colors B. PCA of scRNA-seq data after regressing out the cell cycle phases overlaid with cell cycle phases detected using Cyclone as indicated by the colors C. PCA of scRNA-seq data after regressing out the cell cycle phases overlaid with experimental time points as indicated by the colors D. Distribution of pseudotime values of experimental time points after regressing out the cell cycle phases as in Fig S2C E. Differential gene expression between naive (ESC) and formative pluripotency (Epiblast like cells (EpiLCs), 48 h) obtained by bulk RNA seq F. Venn diagram indicating the number of genes that overlap between the top 500 highly variable genes (HVG) identified by scRNA-seq data (left) and top 500 differentially expressed genes (DEG) in bulk RNA seq data (right) G. Distribution of pseudotime values of scRNA-seq data (as in Fig 1C) at each experimental time point as indicated, plotted as a density plot

**Fig S3 (related to Fig 3):**
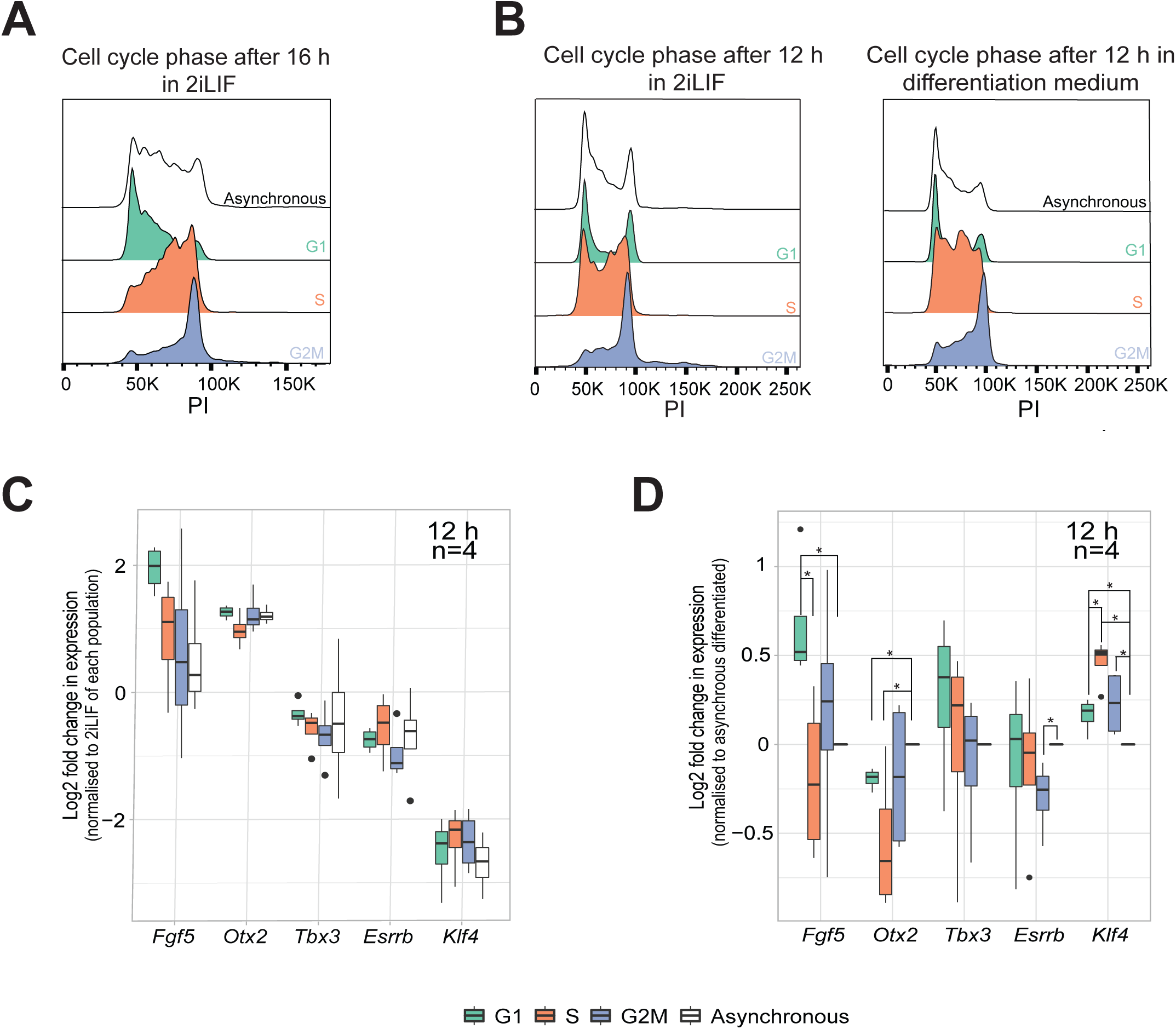
A. Analysis of cell cycle distribution using PI staining, at 16 h post sorting of G1, S, G2M, and asynchronous control sorted cells cultured in 2iLIF B. Analysis of cell cycle distribution using PI staining, at 12 h post sorting of G1, S, G2M, and asynchronous control sorted cells cultured either in 2iLIF (left) or in differentiation medium (right) C. Expression analysis using RT-qPCR of formative *(Fgf5 and Otx2)* and naive markers *(Tbx3, Esrrb, Klf4)* at 12 h post sorting and differentiation. Fold changes of gene expression were calculated by normalization to undifferentiated counterparts of each sorted population and Iog2 transformed. Unpaired Wilcoxon test was performed on the Iog2 transformed values, p<O.05 is considered signifi­cant, n=4 biological replicates D. Expression analysis using RT-qPCR of formative *(Fgf5 and Otx2)* and naive markers *(Tbx3, Esrrb, Klf4)* at 12 h post sorting and differentiation. Fold changes of gene expression were calculated by normalization to the asynchronous differentiated sample and Iog2 transformed. Unpaired Wilcoxon test was performed on the Iog2 transformed values, p<O.05 is considered significant, n=4 biological replicates

**Fig S4 (related to Fig 4):**
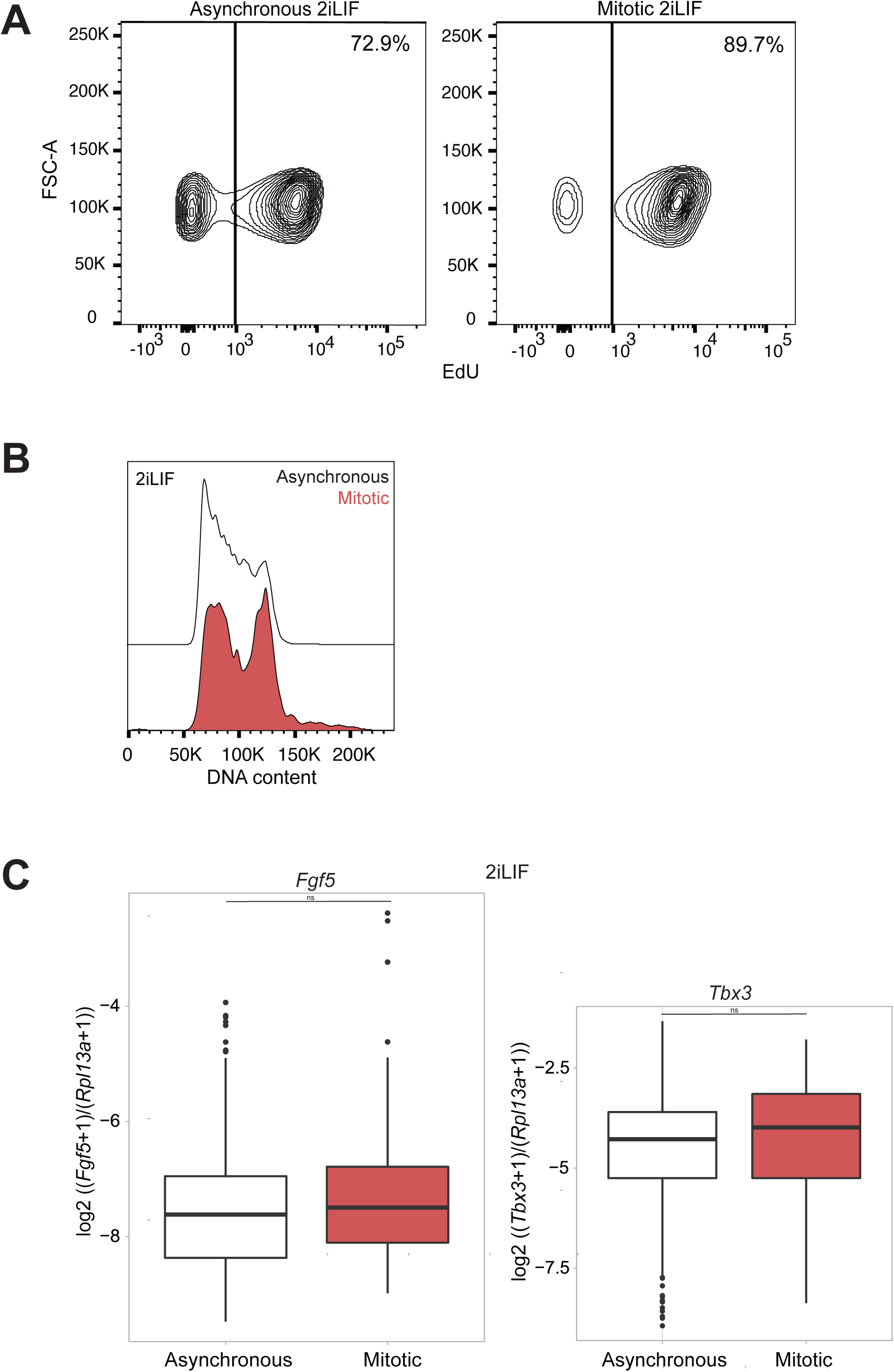
A. Incorporation of EdU measured at 16 h in 2iLIF of asynchronous population (left) and cells released from mitosis (right). 30 mins EdU pulse was administered at 3 h post release from shake-off and 13 h later (total 16 h post release) EdU was developed and the percentage of positive cells was analyzed by FACS (as shown in the schematic of Fig 4C, top). The percentage of positive cells is indicated. B. Distribution of cell cycle at 16 h in 2iLIF after release from mitosis (mitotic) and asynchronous populations, analyzed using Hoechst 33342 C. *Rpl13a* normalized counts of *Fgf5* (left) and *Tbx3* (right) per cell for the asyn­chronous population and samples released from mitosis (mitotic) after 16 h in 2iLIF. calculated as log2 [(no. of *Fgf5* or *Tbx3* transcripts per cell +1) / (no. of *Rpl13a* tran­scripts per cell +1)]. F-test was performed on the log2 transformed values to test the significance of variability between asynchronous population and cells released from mitosis(mitotic). p<0.05 is considered significant. ns: non-significant. One experiment is shown (for the rest of the replicates see S6 I-L)

**Fig S5 (related to Fig 4):**
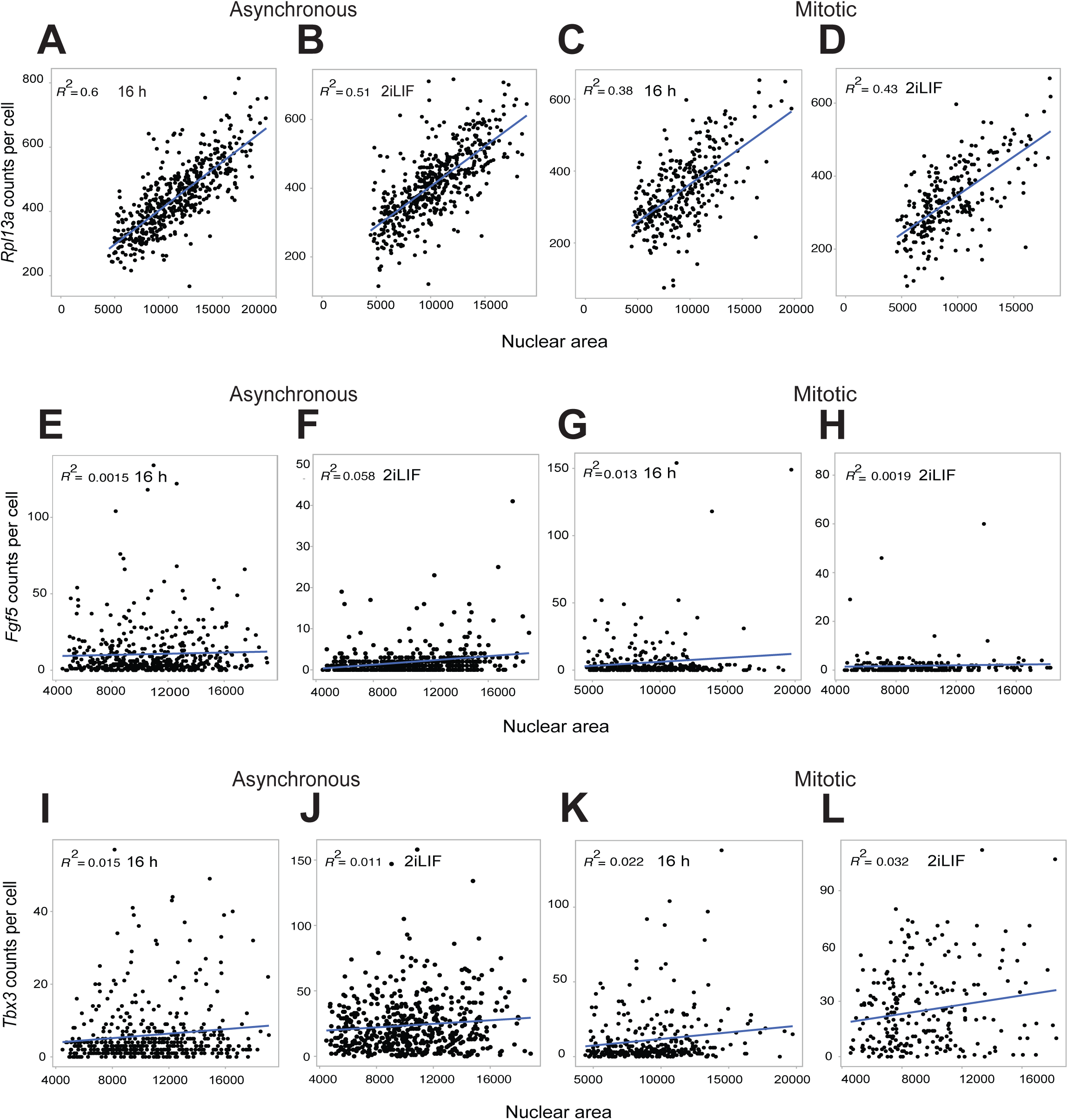
A. Nuclear area vs. number of *Rpl13a* transcripts per cell in asynchronous popula­tion differentiated for 16 h B. Nuclear area vs. number of *Rpl13a* transcripts per cell in asynchronous popula­tion cultured in 2iLIF for 16 h C. Nuclear area vs. number of *Rpl13a* transcripts per cell in cells released from mitosis differentiated for 16 h D. Nuclear area vs. number of *Rpl13a* transcripts per cell in cells released from mitosis cultured in 2iLIF for 16 h E. Nuclear area vs. number of *Fgf5* transcripts per cell in asynchronous population differentiated for 16 h F. Nuclear area vs. number of *Fgf5* transcripts per cell in asynchronous population cultured in 2iLIF for 16 h G. Nuclear area vs. number of *Fgf5* transcripts per cell cells released from mitosis differentiated for 16 h H. Nuclear area vs. number of *Fgf5* transcripts per cell in cells released from mitosis cultured in 2iLIF for 16 h I. Nuclear area vs. number of *Tbx3* transcripts per cell in asynchronous population differentiated for 16 h J. Nuclear area vs. number of *Tbx3* transcripts per cell in asynchronous population cultured in 2iLIF for 16 h K. Nuclear area vs. number of *Tbx3* transcripts per cell in cells released from mitosis differentiated for 16 h L. Nuclear area vs. number of *Tbx3* transcripts per cell in cells released from mitosis cultured in 2iLIF for 16 h One replicate is shown. R^2^ corresponds to the fit of the regression line (shown in blue) plotted using ggplot2 in R.

**Fig S6 (related to Fig 4):**
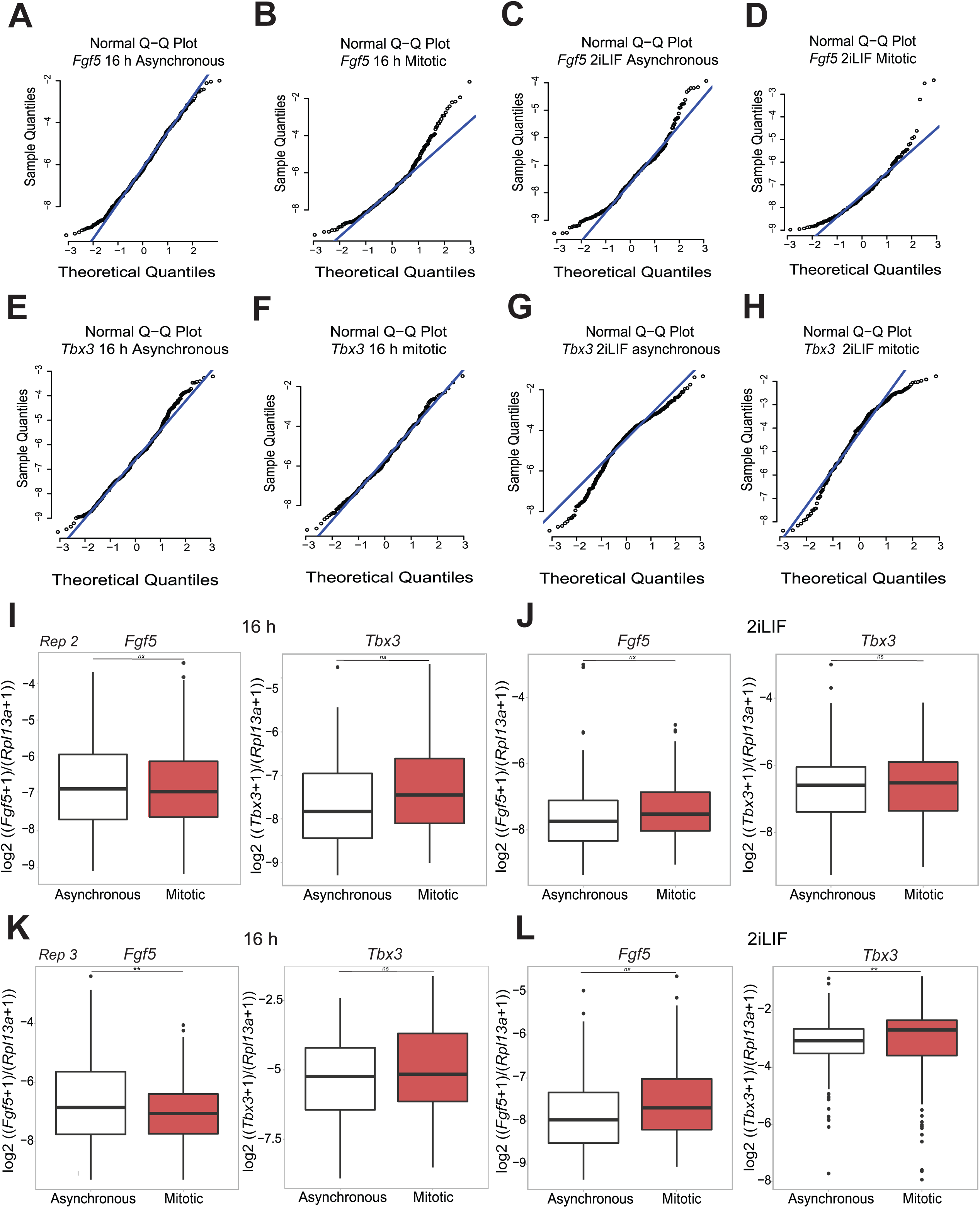
QQ plots of *Rpl13a* normalized counts of each gene per cell calculated as log2 [(no. of transcripts per cell of gene +1)/ (no. of *Rpl13a* per cell +1)] to test the normality of distribution of: A. *Fgf5* at 16 h of asynchronous population in differentiation medium B. *Fgf5* at 16 h post mitotic release in differentiation medium C. *Fgf5* at 16 h of asynchronous populations in 2iLIF D. *Fgf5* at 16 h post mitotic release in 2iLIF E. *Tbx3* at 16 h of asynchronous population in differentiation medium F. *Tbx3* at 16 h post mitotic release in differentiation medium G. *Tbx3* at 16 h of asynchronous population in 2iLIF H. *Tbx3* at 16 h post mitotic release in 2iLIF I. *Rpl13a* normalized counts of *Fgf5* (left) and *Tbx3* (right) per cell for the asyn­chronous population and cells released from mitosis (mitotic) after 16 h of differentia­tion. Calculated as log2 [(no. of *Fgf5* or *Tbx3* transcripts per cell +1) / (no. of *Rpl13a* transcripts per cell +1)]. F-test was performed on the log2 transformed values to test the significance of variability between asynchronous population and cells released from mitosis (mitotic). p<0.05 is considered significant. ns: non-significant. One experiment is shown. *Rep2*- Replicate 2 J. *Rpl13a* normalized counts of *Fgf5* (left) and *Tbx3* (right) per cell for the asyn­chronous population and cells released from mitosis (mitotic) after 16 h in 2iLIF. Calcu­lated as log2 [(no. of *Fgf5* or *Tbx3* transcripts per cell +1) / (no. of *Rpl13a* transcripts per cell +1)]. F-test was performed on the log2 transformed values to test the significance of variability between asynchronous population and cells released from mitosis (mitotic). p<0.05 is considered significant. ns: non-significant. One experiment is shown. *Rep2*-Replicate 2 K. *Rpl13a* normalized counts of *Fgf5* (left) and *Tbx3* (right) per cell for the asyn­chronous population and cells released from mitosis (mitotic) after 16 h of differentia­tion. Calculated as log2 [(no. of *Fgf5* or *Tbx3* transcripts per cell +1) / (no. of *Rpl13a* transcripts per cell +1)]. F-test was performed on the log2 transformed values to test the significance of variability between asynchronous population and cells released from mitosis (mitotic). p<0.05 is considered significant. ns: non-significant. One experiment is shown. *Rep3*- Replicate 3 L. *Rpl13a* normalized counts of *Fgf5* (left) and *Tbx3* (right) per cell for the asyn­chronous population and cells released from mitosis (mitotic) after 16 h in 2iLIF. Calcu­lated as log2 [(no. of *Fgf5* or *Tbx3* transcripts per cell +1) / (no. of *Rpl13a* transcripts per cell +1)]. F-test was performed on the log2 transformed values to test the significance of variability between asynchronous population and cells released from mitosis (mitotic). p<0.05 is considered significant. ns: non-significant. One experiment is shown. *Rep3*-Replicate 3

**Fig S7 (related to Fig 5):**
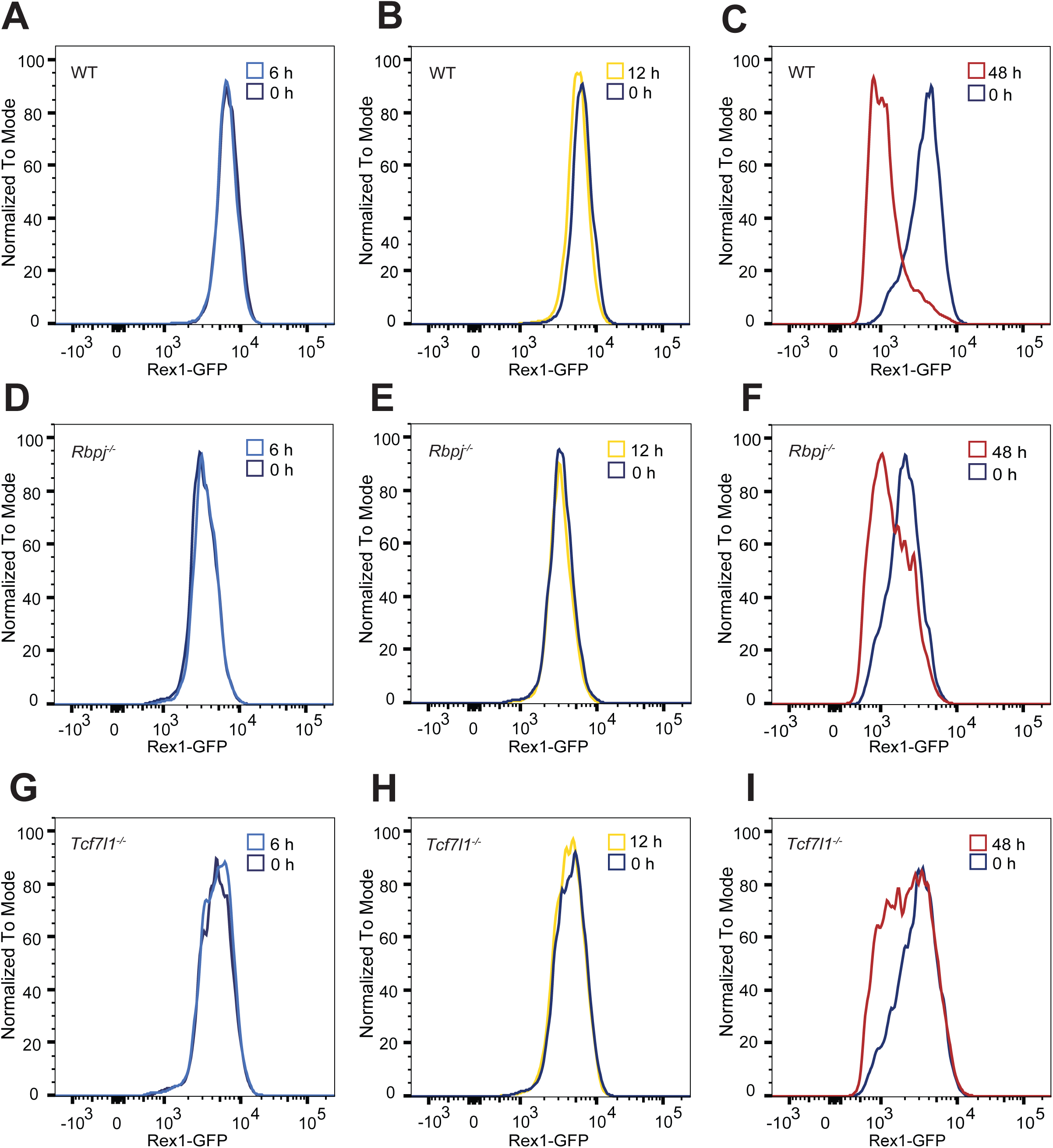
A, D, G: Differentiation status measured by Rex1-GFP levels using FACS of WT (Rex1-GFPd2) (A), *Rbpj^-/-^* (D), *Tcf7l1^-/-^* (G) cultured in 2iLIF (0 h) or 6 h of differentiation B, E, H: Differentiation status measured by Rex1-GFP levels using FACS of WT (Rex1-GFPd2) (B), *Rbpj^-/-^* (E), *Tcf7l1^-/-^* (H) cultured in 2iLIF (0 h) or 12 h of differentiation C, F, I: Differentiation status measured by Rex1-GFP levels using FACS of WT (Rex1-GFPd2) (C), *Rbpj^-/-^* (F), *Tcf7l1^-/-^* (I) cultured in 2iLIF (0 h) or 48 h of differentiation

**Fig S8 (related to Fig 5):**
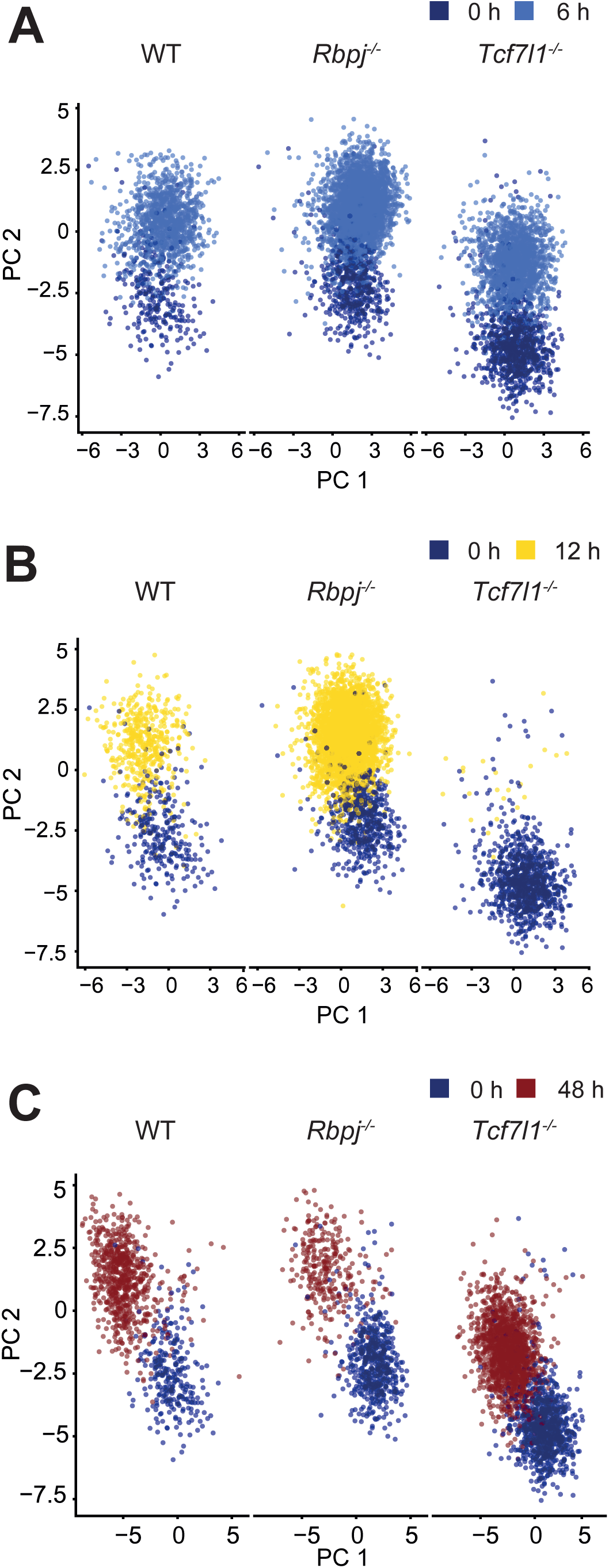
Supervised dimension reduction using principal component analysis (PCA) of WT (Rex1-GFPd2), *Rbpj^-/-^*, and *Tcf7l1^-/-^* cultured in 2iLIF (0 h) or differentiated for: A. 6 h B. 12 h (*Tcf7l1^-/-^* cells were lost during cell pooling for 10X Genomics scRNA-seq) C. 48 h

## REFERENCES

1. Smith, A. Formative pluripotency: the executive phase in a developmental continuum. Dev. Camb. Engl. 144, 365–373 (2017).

2. Nichols, J. & Smith, A. Naive and Primed Pluripotent States. Cell Stem Cell 4, 487–492 (2009).

3. Weinberger, L., Ayyash, M., Novershtern, N. & Hanna, J. H. Dynamic stem cell states: naive to primed pluripotency in rodents and humans. Nat. Rev. Mol. Cell Biol. 17, 155–169 (2016).

4. Wray, J., Kalkan, T. & Smith, A. G. The ground state of pluripotency. Biochem. Soc. Trans. 38, 1027–1032 (2010).

5. Silva, J. et al. Promotion of Reprogramming to Ground State Pluripotency by Signal Inhibition. PLOS Biol. 6, e253 (2008).

6. Ying, Q.-L. et al. The ground state of embryonic stem cell self-renewal. Nature 453, 519–523 (2008).

7. Kinoshita, M. & Smith, A. Pluripotency Deconstructed. Dev. Growth Differ. 60, 44–52 (2018).

8. Pera, M. F. & Rossant, J. The exploration of pluripotency space: Charting cell state transitions in peri-implantation development. Cell Stem Cell 28, 1896–1906 (2021).

9. Morgani, S., Nichols, J. & Hadjantonakis, A.-K. The many faces of Pluripotency: in vitro adaptations of a continuum of in vivo states. BMC Dev. Biol. 17, 7 (2017).

10. Endoh, M. & Niwa, H. Stepwise pluripotency transitions in mouse stem cells. EMBO Rep. 23, e55010 (2022).

11. Buecker, C. et al. Reorganization of Enhancer Patterns in Transition from Naive to Primed Pluripotency. Cell Stem Cell 14, 838–853 (2014).

12. Lackner, A. et al. Cooperative genetic networks drive embryonic stem cell transition from naïve to formative pluripotency. EMBO J. 40, e105776 (2021).

13. Kalkan, T. et al. Complementary Activity of ETV5, RBPJ, and TCF3 Drives Formative Transition from Naive Pluripotency. Cell Stem Cell 24, 785–801.e7 (2019).

14. Mulas, C., Kalkan, T. & Smith, A. NODAL Secures Pluripotency upon Embryonic Stem Cell Progression from the Ground State. Stem Cell Rep. 9, 77–91 (2017).

15. Carbognin, E. et al. Esrrb guides naive pluripotent cells through the formative transcriptional programme. Nat. Cell Biol. 25, 643–657 (2023).

16. Kalkan, T. et al. Tracking the embryonic stem cell transition from ground state pluripotency. Development 144, 1221–1234 (2017).

17. Strawbridge, S. E., Blanchard, G. B., Smith, A., Kugler, H. & Martello, G. Embryonic stem cells commit to differentiation by symmetric divisions following a variable lag period. 10.1101/2020.06.17.157578 (2020) doi:10.1101/2020.06.17.157578.

18. Kolodziejczyk, A. A. et al. Single Cell RNA-Sequencing of Pluripotent States Unlocks Modular Transcriptional Variation. Cell Stem Cell 17, 471–485 (2015).

19. Kumar, R. M. et al. Deconstructing transcriptional heterogeneity in pluripotent stem cells. Nature 516, 56–61 (2014).

20. Toyooka, Y., Shimosato, D., Murakami, K., Takahashi, K. & Niwa, H. Identification and characterization of subpopulations in undifferentiated ES cell culture. Development 135, 909– 918 (2008).

21. Guo, G. et al. Serum-Based Culture Conditions Provoke Gene Expression Variability in Mouse Embryonic Stem Cells as Revealed by Single-Cell Analysis. Cell Rep. 14, 956–965 (2016).

22. Liu, L., Michowski, W., Kolodziejczyk, A. & Sicinski, P. The cell cycle in stem cell proliferation, pluripotency and differentiation. Nat. Cell Biol. 21, 1060–1067 (2019).

23. Soufi, A. & Dalton, S. Cycling through developmental decisions: how cell cycle dynamics control pluripotency, differentiation and reprogramming. Development 143, 4301–4311 (2016).

24. Dalton, S. & Coverdell, P. D. Linking the cell cycle to cell fate decisions. Trends Cell Biol. 25, 592–600 (2015).

25. Zaveri, L. & Dhawan, J. Cycling to Meet Fate: Connecting Pluripotency to the Cell Cycle. Front. Cell Dev. Biol. 6, 57 (2018).

26. Pauklin, S. & Vallier, L. The Cell-Cycle State of Stem Cells Determines Cell Fate Propensity. Cell 155, 135–147 (2013).

27. Singh, A. M. et al. Cell-Cycle Control of Developmentally Regulated Transcription Factors Accounts for Heterogeneity in Human Pluripotent Cells. Stem Cell Rep. 1, 532–544 (2013).

28. Singh, A. M. et al. Cell-Cycle Control of Bivalent Epigenetic Domains Regulates the Exit from Pluripotency. Stem Cell Rep. 5, 323–336 (2015).

29. Pelham-Webb, B. et al. H3K27ac bookmarking promotes rapid post-mitotic activation of the pluripotent stem cell program without impacting 3D chromatin reorganization. Mol. Cell 81, 1732–1748.e8 (2021).

30. Asenjo, H. G. et al. Polycomb regulation is coupled to cell cycle transition in pluripotent stem cells. Sci. Adv. 6, eaay4768 (2020).

31. Waisman, A. et al. Inhibition of Cell Division and DNA Replication Impair Mouse-Naïve Pluripotency Exit. J. Mol. Biol. 429, 2802–2815 (2017).

32. Wray, J. et al. Inhibition of glycogen synthase kinase-3 alleviates Tcf3 repression of the pluripotency network and increases embryonic stem cell resistance to differentiation. Nat. Cell Biol. 13, 838–845 (2011).

33. Chaigne, A. et al. Abscission Couples Cell Division to Embryonic Stem Cell Fate. Dev. Cell 55, 195–208.e5 (2020).

34. McGinnis, C. S. et al. MULTI-seq: sample multiplexing for single-cell RNA sequencing using lipid-tagged indices. Nat. Methods 16, 619–626 (2019).

35. Scialdone, A. et al. Computational assignment of cell-cycle stage from single-cell transcriptome data. Methods 85, 54–61 (2015).

36. Waisman, A. et al. Cell cycle dynamics of mouse embryonic stem cells in the ground state and during transition to formative pluripotency. Sci. Rep. 9, 8051 (2019).

37. Padovan-Merhar, O. et al. Single Mammalian Cells Compensate for Differences in Cellular Volume and DNA Copy Number through Independent Global Transcriptional Mechanisms. Mol. Cell 58, 339–352 (2015).

38. Boileau, R. M., Chen, K. X. & Blelloch, R. Loss of MLL3/4 decouples enhancer H3K4 monomethylation, H3K27 acetylation, and gene activation during embryonic stem cell differentiation. Genome Biol. 24, 41 (2023).

39. Davidson, K. C., Mason, E. A. & Pera, M. F. The pluripotent state in mouse and human. Development 142, 3090–3099 (2015).

40. Guo, G. et al. Klf4 reverts developmentally programmed restriction of ground state pluripotency. Development 136, 1063–1069 (2009).

41. Neganova, I., Zhang, X., Atkinson, S. & Lako, M. Expression and functional analysis of G1 to S regulatory components reveals an important role for CDK2 in cell cycle regulation in human embryonic stem cells. Oncogene 28, 20–30 (2009).

42. Stead, E. et al. Pluripotent cell division cycles are driven by ectopic Cdk2, cyclin A/E and E2F activities. Oncogene 21, 8320–8333 (2002).

43. Huurne, M. ter, Chappell, J., Dalton, S. & Stunnenberg, H. G. Distinct Cell-Cycle Control in Two Different States of Mouse Pluripotency. Cell Stem Cell 21, 449–455.e4 (2017).

44. Jääger, K., et al. G2M splits mouse embryonic stem cells into naïve and formative pluripotency states. 10.1101/616516 (2019) doi:10.1101/616516.

45. Levy, S. H. et al. Esrrb is a cell-cycle-dependent associated factor balancing pluripotency and XEN differentiation. Stem Cell Rep. 17, 1334–1350 (2022).

46. Bunt, J. et al. OTX2 directly activates cell cycle genes and inhibits differentiation in medulloblastoma cells. Int. J. Cancer 131, E21–32 (2012).

47. Trimarchi, J. M., Stadler, M. B. & Cepko, C. L. Individual Retinal Progenitor Cells Display Extensive Heterogeneity of Gene Expression. PLOS ONE 3, e1588 (2008).

48. Raj, A. & van Oudenaarden, A. Nature, Nurture, or Chance: Stochastic Gene Expression and Its Consequences. Cell 135, 216–226 (2008).

49. Rodriguez, J. et al. Intrinsic Dynamics of a Human Gene Reveal the Basis of Expression Heterogeneity. Cell 176, 213–226.e18 (2019).

50. Nair, G., Walton, T., Murray, J. I. & Raj, A. Gene transcription is coordinated with, but not dependent on, cell divisions during C. elegans embryonic fate specification. Development 140, 3385–3394 (2013).

51. Habib, S. J. et al. A Localized Wnt Signal Orients Asymmetric Stem Cell Division in Vitro. Science 339, 1445–1448 (2013).

52. Deathridge, J., Antolović, V., Parsons, M. & Chubb, J. R. Live imaging of ERK signalling dynamics in differentiating mouse embryonic stem cells. Development 146, dev172940 (2019).

53. Aranda, S. et al. Thymine DNA glycosylase regulates cell-cycle-driven p53 transcriptional control in pluripotent cells. Mol. Cell 0, (2023).

54. Kukreja, K., Patel, N., Megason, S. G. & Klein, A. M. Global decoupling of cell differentiation from cell division in early embryo development. 10.1101/2023.07.29.551123 (2023) doi:10.1101/2023.07.29.551123.

55. Nair, G., Walton, T., Murray, J. I. & Raj, A. Gene transcription is coordinated with, but not dependent on, cell divisions during C. elegans embryonic fate specification. Development 140, 3385–3394 (2013).

56. Leyes Porello, E. A., Trudeau, R. T. & Lim, B. Transcriptional bursting: stochasticity in deterministic development. Development 150, dev201546 (2023).

57. Yang, X. et al. Silent transcription intervals and translational bursting lead to diverse phenotypic switching. Phys. Chem. Chem. Phys. 24, 26600–26608 (2022).

58. Jeziorska, D. M. et al. On-microscope staging of live cells reveals changes in the dynamics of transcriptional bursting during differentiation. Nat. Commun. 13, 6641 (2022).

59. Fritzsch, C. et al. Estrogen-dependent control and cell-to-cell variability of transcriptional bursting. Mol. Syst. Biol. 14, e7678 (2018).

60. Koressaar, T. & Remm, M. Enhancements and modifications of primer design program Primer3. Bioinformatics 23, 1289–1291 (2007).

