## supplement table primer for "The asynchrony in the exit from naive pluripotency cannot be explained by differences in the cell cycle phase"

| **Primers for qPCR** | **Forward (5' to 3')** | **Reverse (5' to 3')** |
| --- | --- | --- |
| *Fgf5* | GGGATTGTAGGAATACGAGGAGTT | TGGCACTTGCATGGAGTTT |
| *Otx2* | CGACGTTCTGGAAGCTCTGT | TGGCGGCACTTAGCTCTT |
| *Esrrb* | TACCTGAACCTGCCGATTTC | CCCAGTTGATGAGGAACACA |
| *Klf4* | AAGAACAGCCACCCACACTT | GGTAAGGTTTCTCGCCTGTG |
| *Tbx3* | GCATCCTCTCCTGCTGTCTC | GCCGTAGTGGTGGAAATCTT |
| *Rpl13a* | ACAGCCACTCTGGAGGAGAA | AGGCATGAGGCAAACAGTCT |
